# A blue light receptor CRY1 regulates de novo shoot regeneration

**DOI:** 10.64898/2025.12.09.693185

**Authors:** Min Li, Hikaru Sato, Takuya Sakamoto, Yayoi Inui, Kazunari Yamamoto, Ryosuke Makino, Tomonao Matsushita, Sachihiro Matsunaga

**Affiliations:** The University of Tokyo, Japan; Kanagawa University, Japan; Tokyo University of Science, Japan; Kyoto University, Japan

**Author notes:** **Correspondence:** Hikaru Sato, Sachihiro Matsunaga.

**Keywords:** *Arabidopsis thaliana*, Blue light signaling, Photoresponses, Stem cell, Stress response

## Abstract

Tissue culture enables plant transformation and gene editing, yet inefficient regeneration remains a major bottleneck, and the downstream circuitry linking light perception to tissue culture has not been investigated. Here we demonstrate that the blue-light photoreceptor CRYPTOCHROME 1 (CRY1) enhances *Arabidopsis thaliana* shoot regeneration in a blue light dependent manner. In a two-step root-to-shoot tissue culture system, *cry1* mutant exhibits reduced shoot regeneration. Transcriptomic analyses in the *cry1* mutant reveal that CRY1 suppresses the expression of *AUXIN RESPONSE TRANSCRIPTION FACTOR 3 (ARF3)*. Consistently, *arf3* mutant exhibits enhanced shoot regeneration phenotype. Additionally, genetic and protein interaction assays show that ARF3 acts downstream of CRY1 at both transcriptional and post-translational levels. Furthermore, transcriptomic analyses in the *arf3* mutant show that ARF3 activates salicylic acid (SA)-responsive genes that suppress shoot formation, and exogenous SA inhibits regeneration in *arf3* calli. Integrating RNA-seq with cis-motif analysis identifies stress-responsive ARF3 targets within the CRY1 network. Our results establish a novel CRY1-ARF3 regulatory module that links blue-light perception to hormonal signaling during regeneration, revealing how plants integrate environmental cues with growth-defense trade-offs.

**Synopsis:** The blue-light photoreceptor CRY1 promotes de novo shoot regeneration in *Arabidopsis thaliana* by restraining ARF3-mediated SA signaling. It reveals a fundamental trade-off between defense responses and regenerative growth in plants, coordinated by CRY1-ARF3 module.

- CRY1-mediated blue light signaling enhances de novo shoot regeneration from callus.
- *ARF3* acts downstream of CRY1 and CRY1 physically interacts with ARF3.
- ARF3-activated SA pathway represses shoot regeneration.
- CRY1 enhances shoot regeneration by suppressing ARF3-activated stress responsive genes.

## Introduction

Plant regeneration is a fundamental biological process wherein differentiated somatic cells revert to a pluripotent state, enabling the formation of new tissues and organs (Birnbaum and Sánchez Alvarado, 2008; Sugimoto et al., 2019). This remarkable developmental plasticity is particularly vital for sessile organisms like plants, which cannot escape unfavorable environmental conditions or biotic threats (Xu et al., 2006; Zhou et al., 2019). To adapt and survive, plants have evolved various regenerative strategies, with de novo organogenesis being one of the most critical (Motte et al., 2014; Ikeuchi et al., 2016). This regenerative capacity has been extensively harnessed in plant tissue culture, providing a controlled system to study the molecular mechanisms underlying regeneration (Cheng et al., 2013; Ikeuchi et al., 2019). In tissue culture, a widely adopted two-step protocol facilitates de novo shoot regeneration (Valvekens et al., 1988). Explant tissues are first cultured on an auxin-rich callus-inducing medium (CIM) to induce somatic cells to dedifferentiate into pluripotent callus. Subsequently, the callus is transferred to a cytokinin-rich shoot-inducing medium (SIM) to promote organogenesis of shoot meristems. This method has been instrumental in advancing our understanding of plant developmental biology and has practical applications in agriculture, such as clonal propagation and genetic transformation (Hasnain et al., 2022). Phytohormones such as auxin and cytokinin are well-established drivers of this process, determining the fate of new meristems (Zhang et al., 2017; Zhai and Xu, 2021). However, the influence of external environmental factors, notably light, on regeneration has been relatively less understood (Nameth et al., 2013; Ikeuchi et al., 2016).

Light serves not only as the energy source for photosynthesis but also as a crucial informational signal regulating diverse aspects of plant development (Wei et al., 2023; Zhou et al., 2024; Hao et al., 2025). In *Arabidopsis thaliana* (Arabidopsis), multiple photoreceptors such as cryptochromes (CRY1 and CRY2), phototropins (PHOT1 and PHOT2), and phytochromes (PHYA to PHYE) perceive light of specific wavelengths and initiate downstream signaling cascades (Ahmad and Cashmore, 1993; Briggs and Christie, 2002; Quail, 2002; Liu et al., 2020). These photoreceptor-mediated pathways orchestrate light-dependent developmental responses, including hypocotyl elongation, stomatal development, flowering transition (Wang et al., 2024; Bustamante et al., 2025; Chen et al., 2025). Recent studies have expanded this view by revealing that light signaling also profoundly impacts regenerative capacity. For example, transient exposure of explants to low-fluence red light or a period of darkness after wounding can significantly increase subsequent shoot regeneration frequencies, whereas prolonged low-fluence red light can inhibit the ability to regenerate shoots (Wei et al., 2020). Consistently, light promotes shoot regeneration while suppressing root formation, partly through the activity of the transcription factor ELONGATED HYPOCOTYL 5 (HY5), which modulates meristematic fate during regeneration (Chen et al., 2024). Nevertheless, the detailed molecular networks through which light signaling influences de novo organogenesis remain largely unexplored.

Among the spectrum of light qualities, blue light is especially important given its prominent role in plant development (Wang and Lin, 2020). Blue light is perceived by the cryptochromes, CRY1 and CRY2, which act as photoreceptors to initiate downstream signaling (Yang et al., 2017; Qu et al., 2024). Photoactivated CRYs attenuate the activity of the COP1-SPA E3 ubiquitin ligase complex, leading to stabilization of numerous transcription factors that promote photomorphogenic development (Holtkotte et al., 2017). Under blue light, CRYs accumulate in the nucleus and forms photobodies through liquid-liquid phase separation (LLPS) (Liu et al., 2022). These photobodies are dynamic, membrane-less condensates that function as centers for protein interactions and gene regulation (Liu et al., 2022; Ponnu and Hoecker, 2022; Jiang et al., 2023; Liu et al., 2025a). Despite extensive studies on CRY-mediated signaling in plant development, its regulatory mechanisms during de novo shoot regeneration remain unknown.

Recent research has highlighted a complex interplay between shoot regeneration and stress responses, particularly those mediated by salicylic acid (SA) and jasmonates (JAs) (Park et al., 2019; Hernández-Coronado et al., 2022; Koo et al., 2024). For instance, it has been demonstrated that activation of SA-mediated defense pathways inhibited shoot regenerative processes, suggesting a trade-off where resources are allocated between defense and growth. Hypoxia-induced activation of the RELATED TO APETALA 2.12 (RAP2.12) protein promotes SA biosynthesis, leading to suppressed shoot regeneration (Koo et al., 2024). Conversely, certain stress conditions, such as wounding, can paradoxically enhance callus formation and shoot regenerative efficiency. For example, wounding has been shown to induce the expression of WOUND INDUCED DEDIFFERENTIATION1 (WIND1) that promotes cellular reprogramming and shoot regeneration (Iwase et al., 2017). Thus, depending on the context, different stress signals can either inhibit or facilitate regenerative processes. Despite these insights, the detailed molecular mechanisms orchestrating the balance between regeneration and stress responses are insufficiently understood.

Auxin Response Factors (ARFs) are key transcriptional regulators that mediate auxin-dependent gene expression, orchestrating a wide range of developmental processes including hypocotyl elongation, gynoecium patterning, and lateral root initiation (Cho et al., 2014; Simonini et al., 2017; Liu et al., 2018; Shukla et al., 2019; Kuhn et al., 2020). Phylogenetic analyses classify ARFs into three major classes (A, B and C), with distinct regulatory properties (Mutte et al., 2018). While A-class ARFs typically function as auxin-responsive activators of transcription, B-class ARFs act independently of auxin and can repress A-ARF activity (Lavy et al., 2016; Martin-Arevalillo et al., 2019). The stability of ARF proteins is a critical factor influencing auxin-mediated transcriptional outcomes (Lakehal et al., 2019; Jing et al., 2022). A recent study has identified a conserved degradation motif within the DNA-binding domain of both A- and B-class ARFs across plant species (de Roij et al., 2025). This evidence supports that regulated proteolysis of ARFs is a fundamental aspect of auxin signaling. Several ARFs, including ARF1, ARF6, ARF8, and ARF17, have been shown to undergo proteasome-dependent degradation (Salmon et al., 2008; Lakehal et al., 2019), highlighting the importance of precise control over ARF stability. One proposed degradation mechanism involves intrinsically disordered region (IDR) within ARFs (Jing et al., 2022). A deeper understanding of how degradation is orchestrated across different ARF family members will be essential for clarifying how auxin responses are regulated during development and environmental adaptation.

ARFs are increasingly recognized as points of crosstalk between various hormonal and environmental signals. For instance, (BRASSINOSTEROID INSENSITIVE 2 (BIN2) kinase is a component of the brassinosteroid (BR) signaling pathway. Its interaction with ARF7/19 modulates their properties and constitutes a point of crosstalk between auxin and BR signaling pathway to enhance lateral root organogenesis (Cho et al., 2014). ARFs have also been implicated in plant stress and defense responses, as well as in light signaling contexts, highlighting their broad integrative role. In *Oryza sativa* (rice), ARF3 was recently found to participate in an immune signaling module: a pathogen-induced *A Leaf Expressed and Xoo-induced lncRNA 1* (*ALEX1*), binds to the IDR of ARF3 and modulates ARF3’s phase separation behavior, converting it into a functional state to enhance JA-mediated defense and disease resistance (Lei et al., 2025). Exposure of dark-grown seedlings to light, ETHYLENE RESPONSE FACTOR 72 (ERF72) is translocated to the nucleus, and interacts with ARF6 and BRASSINOZOLE-RESISTANT1 (BZR1) to attenuate the transcriptional regulation of target genes of ARF6 and BZR1 (Liu et al., 2018). As a result, hypocotyl growth is inhibited, and seedlings undergo photomorphogenesis.

ARFs are classified as activators or repressors according to their function to regulate transcription. However, recently more and more studies raised the possibility that ARFs may have a dual role as activators and repressors (Zhang et al., 2018; Cancé et al., 2022). For instance, Arabidopsis ARF3 exhibits dual transcriptional activities, acting either as a repressor or activator depending on tissue context and developmental stage, highlighting its functional complexity (Zhang et al., 2018). Recent reports have emphasized ARF3’s involvement in biotic stress responses in rice, enhancing plant pathogen resistance (Lei et al., 2025). However, the detailed molecular mechanisms by which ARF3 activity and stability are regulated during plant regeneration are unknown.

In this study, we uncovered a novel regulatory mechanism wherein the blue light receptor CRY1 promotes shoot regeneration in Arabidopsis through the inhibition of ARF3-mediated stress responses. Through spatiotemporal analyses of CRY1-GFP and ARF3-GFP during shoot regeneration, we elucidated the dynamic expression patterns of CRY1 and ARF3 throughout the process. Our findings demonstrated that CRY1 modulated ARF3 at both transcriptional and post-translational levels. Moreover, ARF3-activated SA pathway repressed shoot regeneration, and candidate stress-responsive genes were identified to possibly activated by ARF3 downstream of CRY1 signaling. This CRY1-ARF3 module provides insight into how plants integrate environmental cues with hormonal pathways to balance regeneration and defense responses. Understanding this mechanism offers potential applications in agriculture, such as enhancing crop regeneration and pathogen resistance through targeted genetic modifications.

## Results

### CRY1 is required for de novo shoot regeneration from callus

To examine the effect of different photoreceptors on de novo shoot regeneration, we examined the shoot regeneration capacity of knockout mutants of six photoreceptors, including four blue light receptors (CRY1, CRY2, PHOT1 and PHOT2) and two red light receptors (PHYA and PHYB) (Chory, 2010). To quantify the shoot regeneration capacity in *Arabidopsis thaliana*, we performed a two-step tissue culture (Li et al., 2024) for phenotypic analysis of shoot regeneration to quantify the shoot regeneration capacity using *Arabidopsis thaliana*. Among the mutants of six photoreceptors—*phyA-211*, *phyB-5*, *cry1-304*, *cry2-1*, *phot1 and phot2*—only *cry1-304* (Hereafter referred to as *cry1)* exhibited a significant reduction in shoot regeneration comparing with the wild type (WT) (Figs 1A and B). This observation suggested that CRY1-mediated blue light signaling may positively regulate shoot regeneration. In addition, *cry1* exhibited enhanced root regeneration on SIM relative to WT (Fig 1C). Accordingly, it was investigated whether complementation of CRY1 into *cry1* could rescue the phenotype of *cry1*, which displayed suppression of shoot regeneration and enhancement of root regeneration. The result showed that both the suppression of shoot regeneration and the enhancement of root regeneration in *cry1* were partially rescued by CRY1-GFP expression with its native promoter (pro*CRY1*: g*CRY1*-*GFP*) (Figs 1C-E). Taken together, these results suggest that CRY1 has the function of both enhancement of shoot regeneration and suppression of root regeneration.

**Figure 1.**
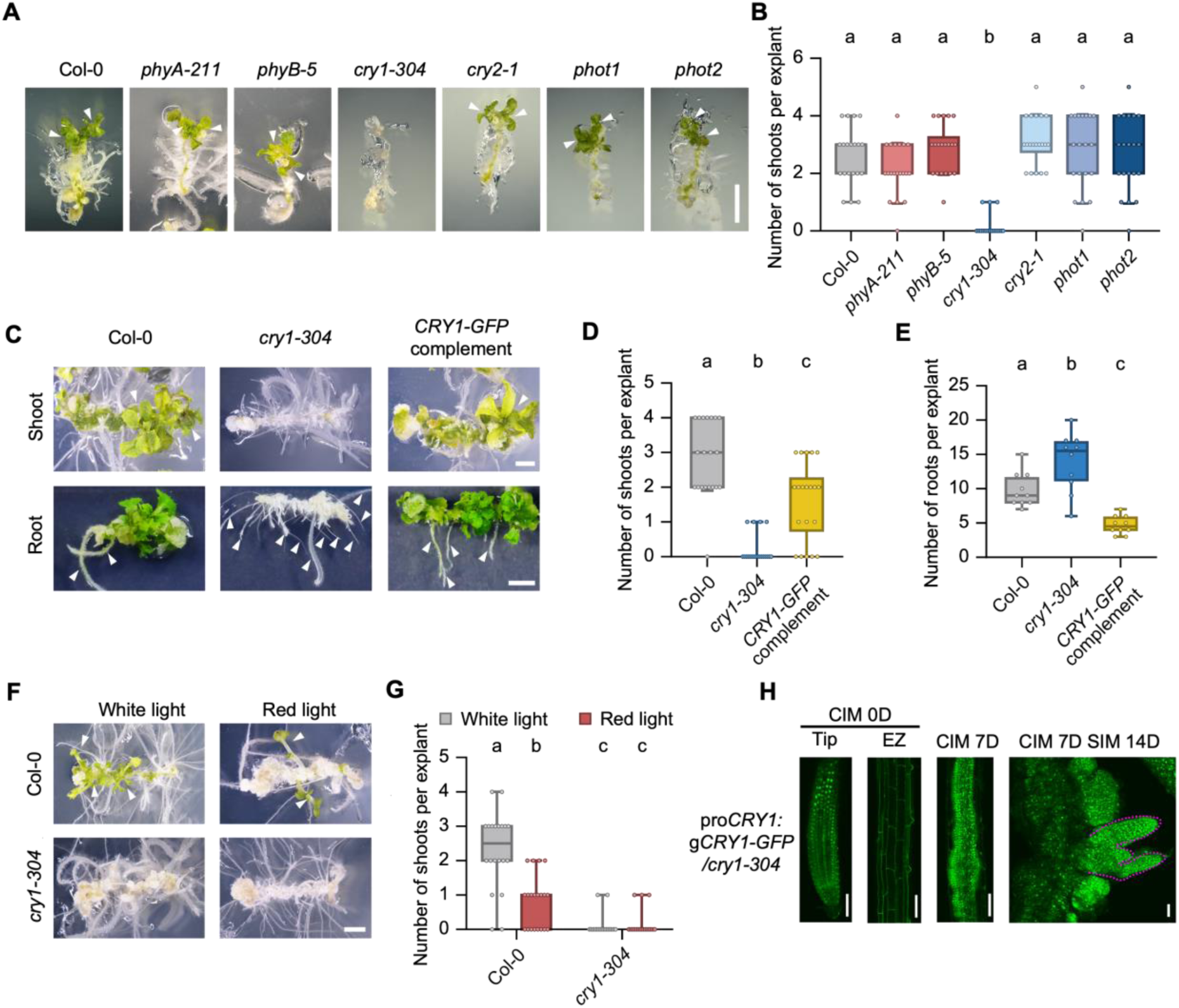
CRY1 is required for de novo shoot regeneration from callus. (A and B) Impact of 6 different photoreceptors on de novo shoot regeneration: the shoot regeneration was reduced only in the *cry1-304* mutant. Shoot regeneration phenotypes of Col-0 and mutants of 6 photoreceptors, *phyA-211*, *phyB-5*, *cry1-304*, *cry2-1*, *phot1* and *phot2*, were observed after root explants were incubated on CIM for 14 days, and then on SIM for 14 days under continuous light (A), and the number of regenerated shoots derived from calli per explant was counted (n = 20) (B). Scale bar: 2 mm. Arrow heads: regenerated shoots. Different letters indicate statistically significant differences, as determined by Tukey’s test (p < 0.05). (C, D and E) CRY1 has both the function to enhance shoot regeneration and to suppress root regeneration. Shoot and root regeneration phenotypes of Col-0, *cry1-304* and the complemented line (pro*CRY1*: g*CRY1-GFP*/*cry1-304*) were observed after root explants were incubated on CIM for 14 days, and then on SIM for 14 days under continuous light (C), and the number of regenerated shoots (D) and roots (E) derived from calli per explant was counted (n = 20). Scale bars: 2 mm. Arrow heads: regenerated shoots/root. Different letters indicate statistically significant differences, as determined by Tukey’s test (p < 0.05). (F and G) CRY1-mediated blue light signaling from callus formation is necessary for shoot regeneration. Shoot regeneration phenotypes of Col-0 and *cry1-304* were observed after root explants were incubated on CIM for 14 days, and then on SIM for 14 days under continuous white light and red light (F), and the number of regenerated shoots derived from calli per explant was counted (n = 20) (G). Scale bar: 2 mm. Arrow heads: regenerated shoots. Different letters indicate statistically significant differences, as determined by Tukey’s test (p < 0.05). (H) Expression pattern of CRY1 in the complemented line, pro*CRY1*: g*CRY1-GFP*/*cry1-304*, throughout shoot regeneration in root explants. Fluorescent signal of CRY1-GFP was observed at root tip (Tip) and elongation zone (EZ) before transferring to CIM, in callus after 7 days on CIM and in regenerating shoot after 14 days on SIM. Scale bars: 50 μm.

Given that the blue light receptor CRY1 is activated by receiving blue light and functions as an initial regulator of signal-transduction factors downstream from it (Ohgishi et al., 2004; Yu et al., 2010), it was investigated whether blocking the blue light would suppress shoot regeneration. Phenotypic analysis of shoot regeneration was conducted in WT and *cry1* under continuous red light and white light from callus induction. The results showed that first, the shoot regeneration capacity of WT significantly decreased under red light, compared to under white light. By contrast, *cry1* exhibited no significant difference in shoot regeneration capacity under red light, compared to under white light. Second, the significant difference in shoot regeneration capacity was observed between WT and *cry1* under red light (Figs 1F and G). These findings suggest that CRY1 contributes to shoot regeneration through both blue light-dependent and -independent mechanisms, although regeneration efficiency is considerably lower without blue light.

Furthermore, since cell-type-specific and developmental-specific expression profiles of CRY1 have not yet been studied throughout shoot regeneration process, we examined the expression pattern of CRY1 during shoot regeneration in root explants using a transgenic complementation line (pro*CRY1*: g*CRY1*-*GFP*/*cry1-304*). Imaging observation (Fig 1H) showed that before callus induction (CIM 0D), fluorescent signal of CRY1-GFP was predominantly localized in the nuclei and was expressed in the root tip including root apical meristem (RAM), while fluorescent signal of CRY1-GFP was not detected in the elongation zone. After transferring to CIM for 7 days (CIM 7D), the fluorescent signal of CRY1-GFP was observed in the entirety of the callus tissue. Subsequently, after transferring to SIM for 14 days following the incubation on CIM for 7 days (CIM 7D SIM 14D), fluorescent signal of CRY1-GFP was observed in the regenerating shoot including both the shoot apical meristem (SAM) and the callus. The expression pattern of CRY1 further supported the hypothesis that CRY1 was required for de novo shoot regeneration from callus.

### Suppression of stress responses and root formation by CRY1 is required for shoot regeneration

To decipher the mechanism by which CRY1 promoted de novo shoot regeneration, RNA sequencing (RNA-seq) was performed in *cry1* and WT using root explants. Root tips (1 cm root explants) excised from seedlings were collected at three distinct stages: 1) before callus induction (C0), 2) 14 days after callus formation (C14), 3) 1 day after shoot formation following 14 days after callus formation (C14S1) (Fig 2A). Transcriptome comparisons revealed differentially expressed genes (DEGs), displaying that 439, 320 and 262 genes were significantly upregulated (FDR < 0.05, Log_2_FC > 1) and 239, 240 and 248 genes were significantly downregulated (FDR < 0.05, Log_2_FC < -1) in *cry1* compared with that in WT at C0, C14 and C14S1 stages, respectively (Fig 2B) (Supplementary Table 2-4).

**Figure 2.**
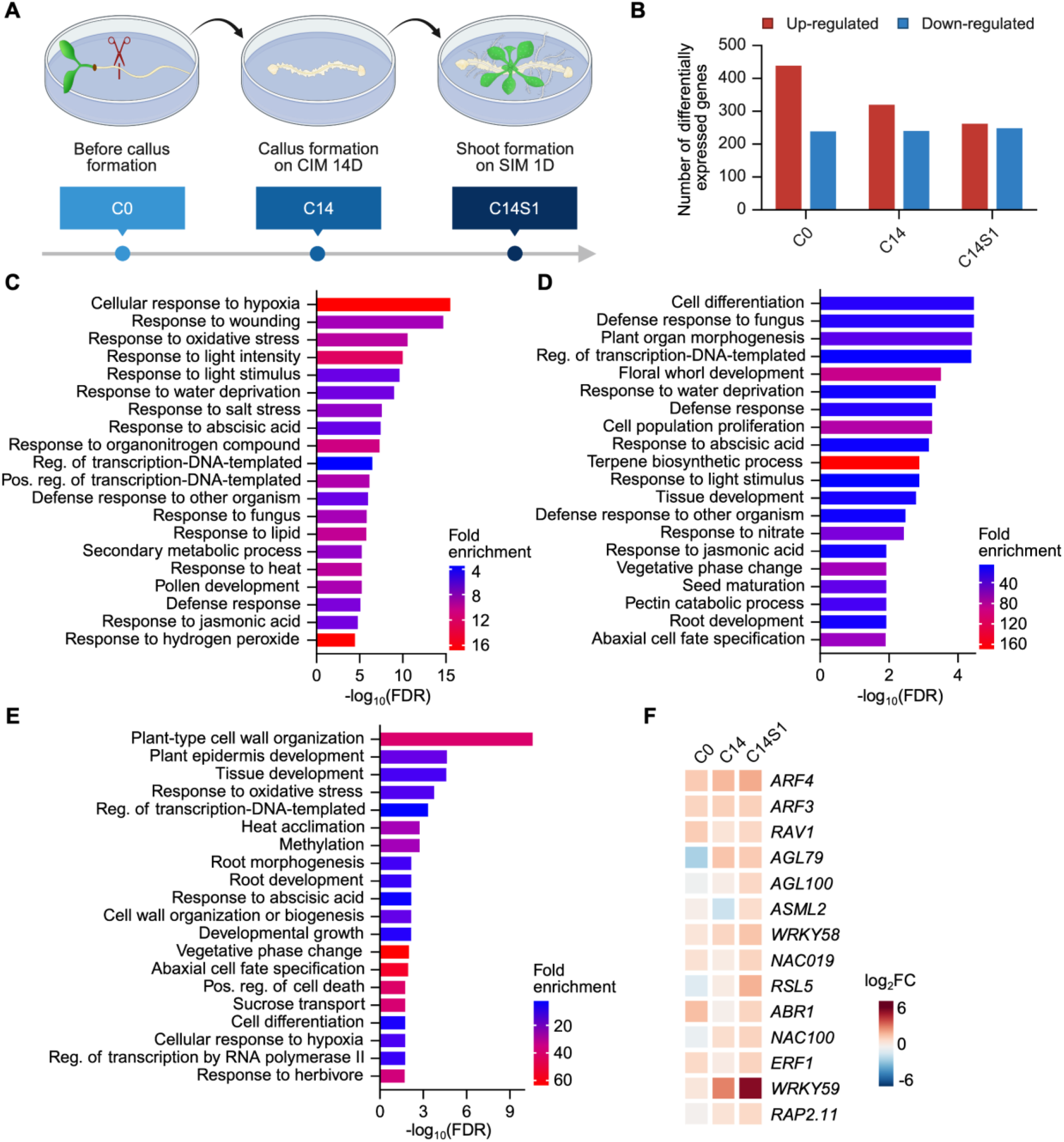
Transcriptomic analysis in *cry1-304* during shoot regeneration. (A) Schematic of the experimental design to study the transcriptomic profile of *cry1-304* during de novo shoot regeneration. The experiment was performed using Col-0 and *cry1-304*. Root explants were collected at three time points: 1) before callus induction (C0), 2) 14 days after callus formation (C14), 3) 1 day after shoot formation following 14 days after callus formation (C14S1). (B) Bar plots indicated the number of up- (FDR < 0.05, Log_2_FC > 1) and down-regulated (FDR < 0.05, Log_2_FC < -1) genes in *cry1-304*. (C, D and E) Top 20 gene ontology (GO) terms enriched in genes up-regulated in *cry1-304* compared with that in WT at C0 (C), C14 (D), and C14S1 (E) stages. (F) Heatmap indicated the overall upregulated patterns of transcription factors with Log_2_FC > 1 in *cry1-304* at C14S1 stage, compared across C0, C14, and C14S1 stages.

Gene ontology (GO) analysis of these DEGs showed that stress responses related biological processes were enriched among upregulated genes in *cry1* compared with that in WT at C0, C14 and C14S1 stages (Figs 2C-E). Moreover, root formation related biological processes were enriched among upregulated genes in *cry1* compared with that in WT at C14 and C14S1 (Figs 2D and E). By contrast, GO analysis showed that photosynthesis-related biological processes were enriched among downregulated genes in *cry1* compared with that in WT at C0, C14 and C14S1 stages (Appendix Fig S1A–C). These results suggest that suppression of stress responses and root formation, along with enhancement of photosynthesis by CRY1 may be critical for shoot regeneration. The evidence of enhanced root regeneration and suppressed shoot regeneration in *cry1* (Fig 1C) supported the hypothesis that suppression of root formation by CRY1 may be critical for shoot regeneration. Previously reported negative role of CRY1-mediated blue-light signaling in plant thermotolerance (Liu et al., 2025b); negative role of CRY1 in lateral rooting (Zeng et al., 2010), as well as positive role of CRY1 in stomatal opening for photosynthesis (Hao et al., 2025) also supported our hypothesis. Moreover, the suppressive effects of CRY1 on stress responses are neither callus- nor shoot-regeneration-specific, while the suppressive effects of CRY1 on root formation are callus- and shoot-regeneration-specific (Figs 2C-E).

### *ARF3* acts downstream from CRY1 to inhibit shoot regeneration

Transcriptomic analysis revealed widespread misregulated expression of various genes in *cry1*. It promoted an investigation into which transcription factors predominantly contributed to the transcriptomic and phenotypic divergence between *cry1* and WT. To uncover the underlying mechanism, we first examined the GO enrichment among upregulated genes in *cry1* at C14S1 (Fig 2E). GO analysis highlighted a significant enrichment of genes associated with root formation and cell wall organization, both of which are processes critically regulated by auxin signaling (Atkinson et al., 2014; Schoenaers et al., 2018). Moreover, phenotypic analysis revealed enhanced root regeneration in *cry1* (Fig 1C), consistent with a higher auxin to cytokinin ratio during root regeneration process. Consequently, the investigation shifted to the 262 upregulated genes in *cry1* at C14S1 with the objective of identifying auxin-related transcription factors. Among these, 14 transcription factors were identified as transcriptional downstream targets of CRY1 (Fig 2F). Notably, *ARF3* and *ARF4*, which are transcription factors in the response to auxin, were selected as major candidate downstream targets of CRY1 due to their well-characterized roles in auxin response (Cheng et al., 2013; Guan et al., 2017).

To evaluate whether the upregulation of *ARF3* or *ARF4* contributes to the suppressed shoot regeneration phenotype in *cry1*, a phenotypic analysis of shoot regeneration was performed in *ett-13/arf3* (Pekker et al., 2005) (hereafter referred to as *arf3)* and *arf4-1* (Pekker et al., 2005). Compared to WT, *arf3* exhibited a significantly higher shoot regeneration, while *arf4-1* displayed a markedly lower shoot regeneration, a phenotype that is consistent with a previous report (Zhang et al., 2021) (Figs 3A and B). These results revealed an opposite shoot regeneration phenotype between *cry1* and *arf3*, but not between *cry1* and *arf4-1*. Based on these observations, we identified *ARF3* as a potential downstream target of CRY1 in the regulation of shoot regeneration.

**Figure 3.**
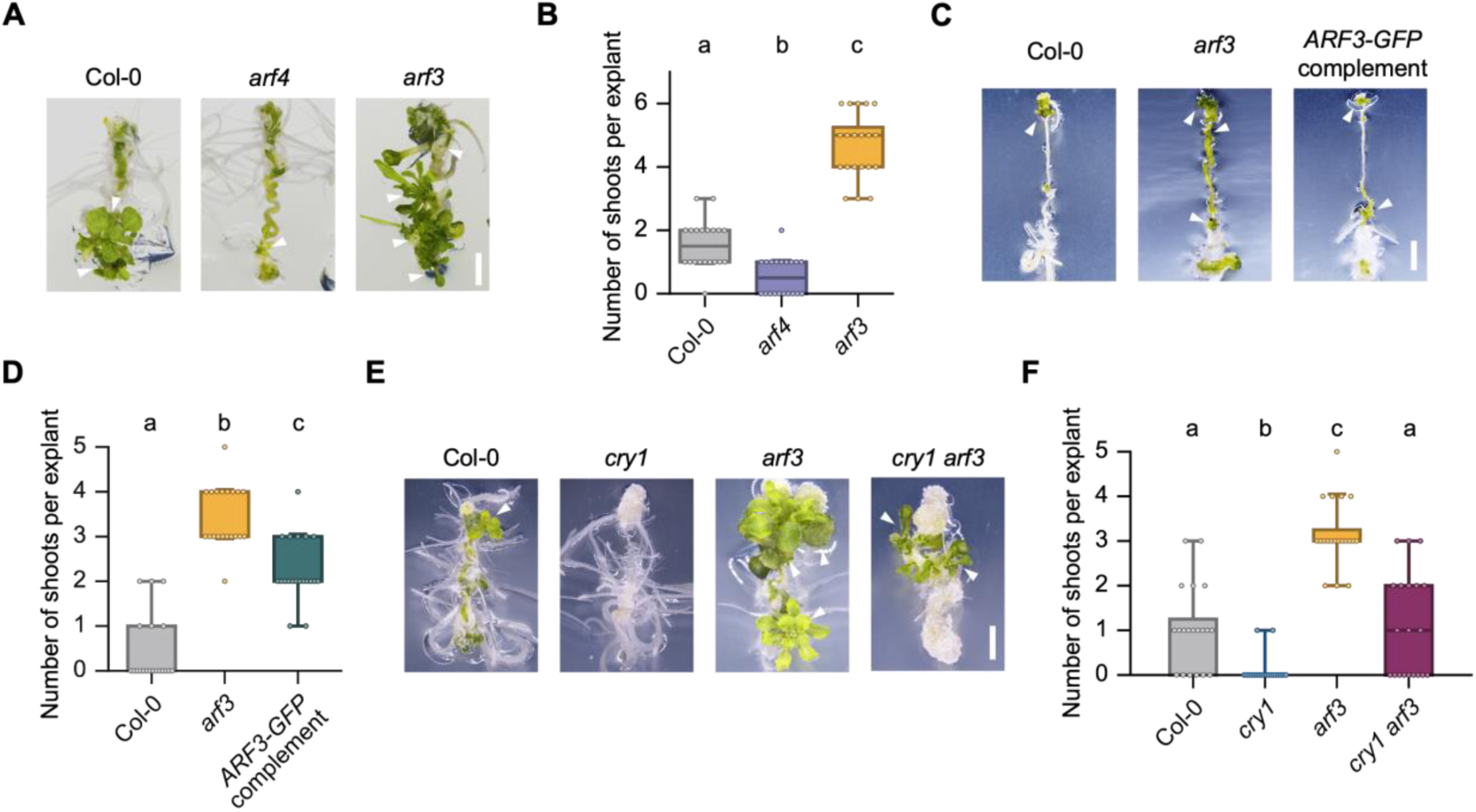
*ARF3* acts downstream from CRY1 to inhibit shoot regeneration. (A and B) ARF3 had negative effects on shoot regeneration. Shoot regeneration phenotypes of Col-0, *arf4* and *arf3* were observed after root explants were incubated on CIM for 14 days, and then on SIM for 14 days under continuous light (A), and the number of regenerated shoots derived from calli per explant was counted (n = 20) (B). Scale bar: 2 mm. Arrow heads: regenerated shoots. Different letters indicate statistically significant differences, as determined by Tukey’s test (p < 0.05). (C and D) ARF3 had the function to repress shoot regeneration. Shoot regeneration phenotypes of Col-0, *arf3* and the complemented line (pro*ARF3*: g*ARF3-GFP*/*arf3*) were observed after root explants were incubated on CIM for 14 days, and then on SIM for 7 days under continuous light (C), and the number of regenerated shoots derived from calli per explant was counted (n = 20) (D). Scale bar: 2 mm. Arrow heads: regenerated shoots. Different letters indicate statistically significant differences, as determined by Tukey’s test (p < 0.05). (E and F) The *cry1 arf3* double mutant rescued the shoot regeneration phenotype of the *cry1* single mutant. Shoot regeneration phenotypes of Col-0, *cry1*, *arf3* and *cry1 arf3* were observed after root explants were incubated on CIM for 14 days, and then on SIM for 14 days under continuous light (E), and the number of regenerated shoots derived from calli per explant was counted (n = 20) (F). Scale bar: 2 mm. Arrow heads: regenerated shoots. Different letters indicate statistically significant differences, as determined by Tukey’s test (p < 0.05).

To further investigate the role of ARF3, we explored whether the complementation of ARF3 into a*rf3* could rescue the phenotype of *arf3*. The results demonstrated that the enhancement of shoot regeneration in *arf3* was partially complemented by ARF3-GFP expression driven by its native promoter (pro*ARF3*: g*ARF3*-*GFP*) (Figs 3C and D), indicating that ARF3 functions as a negative regulator of shoot regeneration. Additionally, to confirm the genetic interaction between *CRY1* and *ARF3,* and to investigate the role of *ARF3* in the downstream of CRY1 in the regulation of shoot regeneration, phenotypic analysis of shoot regeneration was examined in *cry1 arf3* double mutant. The *cry1 arf3* double mutant exhibited significantly higher shoot regeneration compared to the *cry1* single mutant (Fig 3E and F), demonstrating that *ARF3* genetically acts downstream of *CRY1* in repressing shoot regeneration.

### CRY1 physically interacts with ARF3 in a light-independent manner

Because CRY1 is well known for its role in regulating transcriptional activity of transcription factors (Chaves et al., 2011), we examined whether ARF3 is subject to post-translationally regulated by CRY1. Previously, CRY1 has been reported to interact physically with ARF6 and ARF8 in a blue light-dependent manner (Mao et al., 2020). To investigate whether CRY1 also interacts with ARF3 to regulate shoot regeneration, we performed bimolecular fluorescence complementation (BiFC) assays in *Agrobacterium*-infiltrated *Nicotiana benthamiana* (tobacco) leaves. Upon transient expression in tobacco leaves incubated under light for 4 days, strong fluorescence signals were detected in the nuclei of epidermal cells co-transformed with *CRY1-cYFP* and *nYFP-ARF3*, whereas no fluorescence signal was detected in epidermal cells transformed separately with either *CRY1-cYFP* or *nYFP-ARF3* (Fig 4A). These results demonstrated a physical interaction between CRY1 and ARF3 in nuclei.

**Figure 4.**
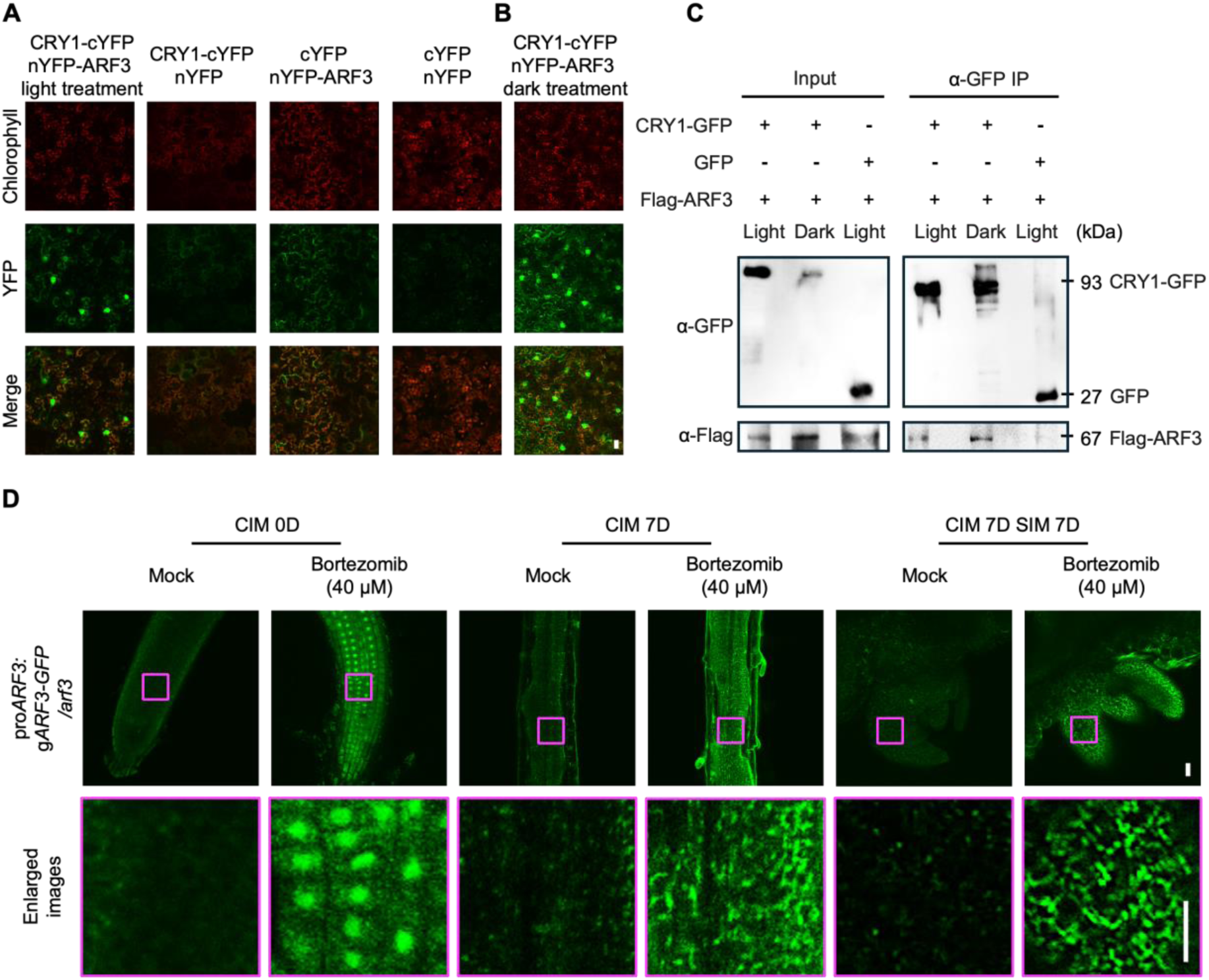
CRY1 physically interacts with ARF3 in a blue light-independent manner. (A and B) Bimolecular fluorescence complementation (BiFC) assays showing the interactions of CRY1 with ARF3 in the epidermal cells of *Nicotiana benthamiana* (tobacco) leaves. The transient expression in tobacco leaves were incubated under light treatment for 4 days (A), and incubated under dark treatment for 1 day, following light treatment for 4 days (B). Confocal images of BiFC fluorescence (YFP), chlorophyll autofluorescence and merged images are shown. Scale bar: 20 μm. (C) Co-immunoprecipitation (Co-IP) assay showing the interactions of CRY1 with ARF3. The GFP-tagged CRY1 or the control GFP were coexpressed with Flag-tagged ARF3 in tobacco leaves, respectively. The transient expression in tobacco leaves were incubated under light and dark condition for 5 days. Total protein extracts (Input) or immunoprecipitated fractions using an anti-GFP antibody (α-GFP IP) were analyzed by immunoblotting using anti-GFP and anti-FLAG antibodies. (D) Expression pattern of ARF3 in the complemented line, pro*ARF3*: g*ARF3-GFP*/*arf3*, throughout shoot regeneration in root explants in the absence (mock, 0.1% DMSO) and presence of the 26S proteasome inhibitor, bortezomib. Fluorescent signal of ARF3-GFP was observed at root tip before transferring to CIM, in calli after 7 days on CIM and in regenerating shoot after 7 days on SIM. Magenta squares at upper panels are enlarged at lower panels. Scale bars: 20 μm.

To investigate whether the CRY1-ARF3 interaction is dependent on light, an additional BiFC assay was performed with transient expression in tobacco leaves incubated under light for 4 days then under dark for 1 day. Strong fluorescence signals were detected both in nucleus and cytoplasm of epidermal cells co-transformed with *CRY1-cYFP* and *nYFP-ARF3* (Fig 4B). These results indicated that CRY1 interacted with ARF3 in a light-independent manner.

To further validate this interaction independent of light, we conducted co-immunoprecipitation (Co-IP) assays using protein extracts from tobacco leaves co-expressing CRY1 tagged with GFP and ARF3 tagged with Flag, incubated under light for 3 days and incubated in the dark for 3 days, respectively. Co-IP assays showed that under both light and dark conditions, CRY1 immunoprecipitated ARF3, but the control protein GFP did not, demonstrating that CRY1 associates with ARF3 irrespective of blue light exposure (Fig 4C). Collectively, these results provide evidence that CRY1 physically interacts with ARF3 in a light-independent manner.

The enhancement of shoot regeneration capacity in *arf3* implied its involvement in regulating shoot regeneration. To further investigate the function of ARF3 during shoot regeneration, we examined its spatial expression pattern throughout shoot regeneration in root explants using a transgenic complementation line (pro*ARF3*: g*ARF3*-*GFP*/*arf3*). Among our observations, we detected ARF3-GFP signals at CIM 0D, CIM 7D, and CIM 7D SIM 7D stages (Fig 4D); although the signals were very weak in the root tip, callus cells, and SAM, the rescued phenotype in the complemented line confirmed that even low levels of ARF3 are sufficient for its molecular function to suppress shoot regeneration. The weak signals of ARF3-GFP raised the possibility that the limited accumulation of ARF3 might be attributable to degradation. Previous studies showed that CRY1 interacted directly with PIF5, and that PIF5 accumulated under low blue light (LBL) conditions compared with under white light condition (Pedmale et al., 2016). Additionally, CRY1 promoted the NPR1-mediated degradation of PIF4 (Zhou et al., 2024). These findings suggest a potential role for CRY1 in mediating the degradation of its interacting transcription factors.

Accordingly, we investigated the effect of the 26S proteasome inhibitor, bortezomib (Xu et al., 2017), on ARF3-GFP expression throughout shoot regeneration. Treatment with 40 µM bortezomib markedly increased GFP fluorescence intensity compared with the mock control at CIM 0D, CIM 7D, and CIM 7D SIM 7D (Fig 4D), implying that the ARF3 protein is degraded throughout shoot regeneration. Moreover, ARF3-GFP signals were observed in nuclei at CIM 0D, while dot-like structures appeared throughout cells at CIM 7D and CIM 7D SIM 7D, consistent with previous observations in the floral meristem and SAM (Zhang et al., 2022). Collectively, these results suggested that CRY1 may promote the degradation of ARF3 through direct interaction, and that even the small amount of non-degraded ARF3 in the nucleus was functionally sufficient to suppress shoot regeneration.

### ARF3-activated salicylic acid pathway represses shoot regeneration

To clarify the role of ARF3 during shoot regeneration, RNA-seq was performed on root explants from *arf3* and WT, using the same approach employed for *cry1* (Fig 2A). Transcriptome comparisons revealed DEGs: specifically, 312, 390 and 288 genes were significantly upregulated (FDR < 0.05, Log_2_FC > 1), and 236, 846 and 866 genes were significantly downregulated (FDR < 0.05, Log_2_FC < -1) in *arf3* compared with that in WT at C0, C14 and C14S1 stages, respectively (Fig 5A) (Supplementary Table 5-7). The finding that a greater number of genes were downregulated than upregulated in *arf3*, specifically at C14 and C14S1 stages, suggested that ARF3 mainly functioned as a transcriptional activator during shoot regeneration. This transcriptional activation activity of ARF3 aligned with previous findings demonstrating that, under high auxin conditions, ARF3 activated ARF3 targets during gynoecium development (Kuhn et al., 2020).

**Figure 5.**
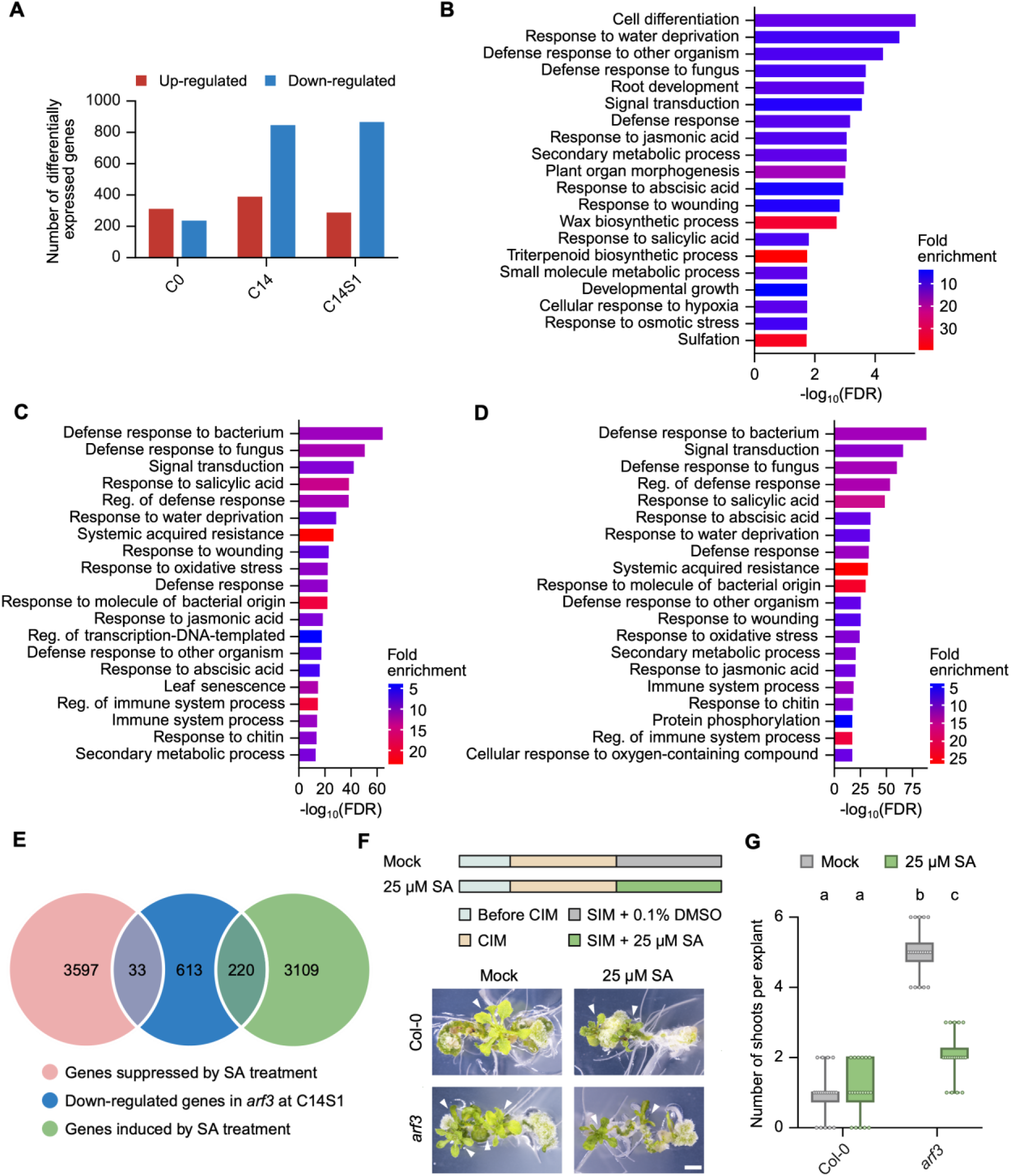
ARF3-activated salicylic acid pathway represses shoot regeneration. (A) Bar plots indicated the number of up- (FDR < 0.05, Log_2_FC > 1) and down-regulated (FDR < 0.05, Log_2_FC < -1) genes in *arf3*. (B, C and D) Top 20 gene ontology (GO) terms enriched in genes down-regulated in *arf3* compared with that in WT at C0 (B), C14 (C), and C14S1 (D) stages. (E) Venn diagrams showing the overlap of the down-regulated genes in *arf3* at C14S1 stage (blue) and the genes suppressed by salicylic acid (SA) treatment (pink) or the genes induced by SA treatment (green) (Zhang et al., 2020). (F and G) *arf3* mutant showed higher sensitivity to SA during shoot regeneration. Shoot regeneration phenotypes of Col-0 and *arf3* were observed after root explants were incubated on CIM for 14 days, and then on SIM in the absence (mock, 0.1% DMSO) and presence of 25 μM SA, for 14 days under continuous light (F), and the number of regenerated shoots derived from calli per explant was counted (n = 20) (G). Scale bar: 2 mm. Arrow heads: regenerated shoots. Different letters indicate statistically significant differences, as determined by Tukey’s test (p < 0.05).

GO analysis of these DEGs revealed that biotic and abiotic stress responses related biological processes were enriched among both upregulated and downregulated genes in *arf3* compared with that in WT at C0, C14 and C14S1 stages (Figs 5B-D; Appendix Fig S2A–C). Given our focus on shoot regeneration rather than callus formation and considering that *cry1* and *arf3* exhibited significantly repressed and enhanced shoot regeneration phenotypes, respectively, we prioritized transcriptomic data from the C14S1 stage. Notably, the ‘response to salicylic acid (SA)’ pathway was enriched among downregulated genes in *arf3* compared with that in WT at C14S1 stage (Fig 5D).

Furthermore, because *SALICYLIC ACID INDUCTION DEFICIENT 2* (*SID2*), a pivotal SA biosynthetic gene reported to repress shoot regeneration (Koo et al., 2024), was among the downregulated genes in *arf3* at the C14S1 stage in our data set (Appendix Fig S2D), we examined the overlap between SA-inducible or SA-suppressive genes (Zhang et al., 2020) and upregulated or downregulated genes in *arf3* at C14S1 stage. Venn diagram analyses revealed a substantial overlap between the downregulated genes in *arf3* at C14S1 stage and genes either induced or suppressed by SA treatment (Fig 5E); notably, the number of overlapping genes was much greater for those induced by SA treatment than for those suppressed by SA treatment. In contrast, the overlaps between the upregulated genes in *arf3* at C14S1 stage and genes either induced or suppressed by SA treatment were both lower than the overlap observed between the downregulated genes in *arf3* and the genes induced by SA treatment (Appendix Fig S2E). Collectively, these analyses indicate that ARF3 positively regulates the expression of salicylic acid-responsive genes during shoot regeneration.

To further validate ARF3’s role in activating stress responses, we noted that a previous study reported that ARF3 upregulated the expression of genes responsive to both biotic and abiotic stresses during early floral development (Zhang et al., 2018). Moreover, ARF3 has been implicated as a key regulator of gynoecium development (Simonini et al., 2017). We then analyzed the GO enrichment of downregulated genes in the *ett-3* loss-of-function mutant compared to WT. Interestingly, biological processes related to biotic and abiotic stress responses were also enriched, with the ‘response to SA’ pathway significantly represented (Appendix Fig S2F), further supporting our claim that ARF3 played a positive role in SA signaling.

Given that current phenotypic analysis revealed that ARF3 inhibited shoot regeneration, and that SA has been reported to negatively affect shoot regeneration (Koo et al., 2024), we next investigated whether this negative effect is mediated by ARF3. We treated WT and *arf3* calli with exogenous SA during shoot induction (Pasternak et al., 2019). In WT, callus cultured on SA-supplemented SIM exhibited no significant difference in shoot regeneration compared with those incubated on normal SIM. By contrast, in *arf3,* callus incubated on SA-supplemented SIM displayed a significantly repressed shoot regeneration capacity compared with the control SIM (Fig 5F and G). These results indicated that WT was insensitive to exogenous SA or at least insensitive to 25 μM SA, possibly because of sufficient endogenous SA levels. In contrast, exogenous SA suppressed shoot regeneration in *arf3* by compensating the repressed SA biosynthesis and activating the signaling pathway. In agreement with the GO enrichment among downregulated gene in *arf3* at C14S1 stage, these findings suggested that ARF3-activated SA pathway repressed shoot regeneration.

### CRY1 enhances shoot regeneration by suppressing ARF3-activated stress responsive genes

ARF proteins have been shown to recognize specific auxin response elements (AuxREs) (Ulmasov et al., 1997), which are typically defined as TGTCNN elements (where N can be any nucleotide) in the promoters of auxin-responsive genes (Guilfoyle and Hagen, 2007; Boer et al., 2014). Published data indicated that ARF3 recognized TGTCGG and TGTCGA as AuxREs based on protein binding microarrays *in vitro* (Franco-Zorrilla et al., 2014), and that ARF3 bound TGTCAT and TGTCAC during gynoecium development (Simonini et al., 2016). To explore potential ARF3-specific AuxREs during shoot regeneration, we searched for cis-acting promoter elements (Maruyama et al., 2012) in the 1 kbp upstream regions of downregulated genes in *arf3* at C14S1 stage. The 20 most-frequent hexamers among all possible hexamer sequences (4^6^ = 4096) were identified (Fig 6A). TGTCAA was recognized as potential ARF3-specific AuxREs during shoot regeneration, as it ranked 17th among the 20 most-frequent hexamers (highlighted in yellow in Fig 6A). The enrichment of the TGTCAA suggested that ARF3 may directly bind these elements in the promoter regions of its target genes, thereby activating their expression during shoot regeneration.

**Figure 6.**
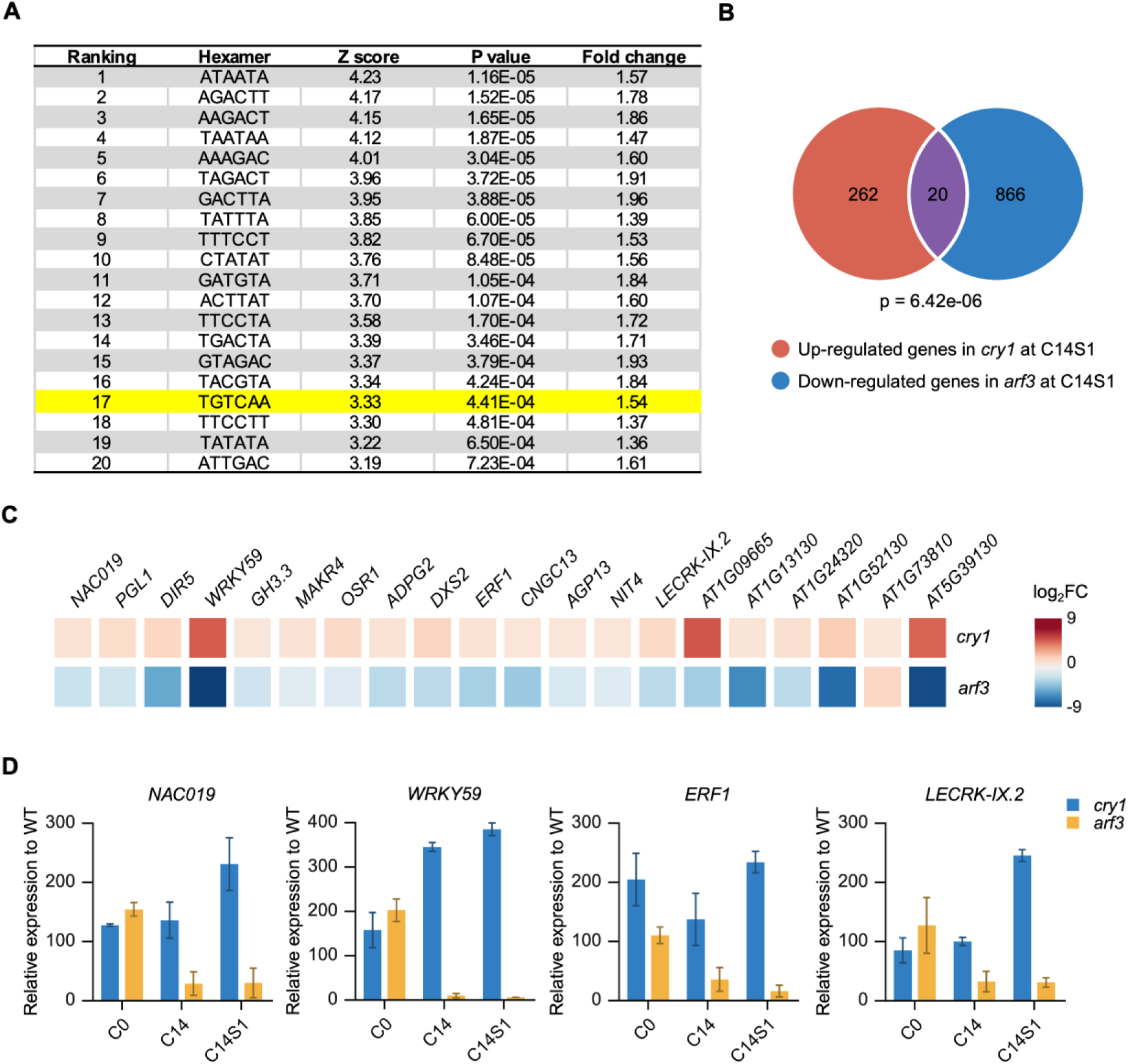
CRY1 enhances shoot regeneration by suppressing ARF3-activated stress responsive genes. (A) The 20 most-frequent hexamers of all possible hexamers (4,096 in total) identified in the promoter regions 1 kb upstream of down-regulated genes in *arf3* at C14S1 stage. The ARF3 specific binding motif, TGTCAA, had the 17th highest Z-score (highlighted in yellow). All *Arabidopsis thaliana* gene promoters were utilized as a control. (B and C) Venn diagram showing the significant overlap of 20 genes (purple) between the up-regulated genes in *cry1* at C14S1 stage (orange) and the down-regulated genes in *arf3* at C14S1 stage (blue) (B). Heatmap illustrating the fold change (log_2_FC) of the 20 overlapping genes (C). (D) Expression patterns of four candidate stress responsive genes, *NAC019*, *WRKY59*, *ERF1* and *LECRK-IX.2*, in *cry1* and *arf3* at C0, C14, and C14S1 stages. The four genes exhibited opposite relative expression patterns in *cry1* and *arf3* at C14S1, respectively. Relative expression was measured as normalized counts per million mapped reads, and the expression level of each gene in WT at C0, C14, C14S1 stage were set to 100.

To identify ARF3-target genes acting downstream of CRY1 during shoot regeneration, we examined the significant overlap (p = 6.42e-06) between downregulated genes in *arf3* and upregulated genes in *cry1* at C14S1 stage, which comprised 20 genes (Fig 6B and C) (Supplementary Table 8). Among these, 13 genes contained potential ARF3-specific AuxREs during shoot regeneration (TGTCAA), and the remaining 7 genes had the core elements of AuxREs (TGTC) in their promoter regions, supporting our analysis of cis-acting promoter elements. Based on the GO enrichment among upregulated genes in *cry1* and downregulated genes in *arf3* at C14S1 stage, we supposed that ARF3-activated stress response pathway may repress shoot regeneration in *cry1*. Consequently, four stress responsive genes, *NAC019*, *WRKY59*, *ERF1* and *LECRK-IX.2* were selected as potential ARF3 target genes in the downstream of CRY1 due to their established roles in stress responses (Berrocal-Lobo et al., 2002; Hu et al., 2013; Luo et al., 2017; Sukiran et al., 2019). Expression patterns of these four stress responsive genes were analyzed at C0, C14 and C14S1 stages using the RNA-seq data in *cry1* and *arf3* (Fig 6D). Relative expression was measured as normalized counts per million mapped reads, and the expression level of each gene in WT at C0, C14 and C14S1 stage were set to 100. Notably, these genes exhibited opposite relative expression patterns in *cry1* and *arf3* at C14S1 stage, respectively. Collectively, these results indicated that ARF3 may directly activate these four stress responsive genes downstream of CRY1 during shoot regeneration.

## Discussion

The present study demonstrated that CRY1-mediated blue light signaling has a critical role in promoting de novo shoot regeneration in *A. thaliana*, acting through the repression of the ARF3-activated SA pathway. CRY1 suppresses ARF3 expression and activity through two distinct mechanisms: first, by downregulating *ARF3* transcription, and second, by means of a direct, light-independent interaction with ARF3. ARF3 expression initially occurs in root tips, subsequently forming dot-like condensates in callus and shoot apical meristem regions. The ARF3 protein undergoes proteasomal degradation during the process of shoot regeneration, whereas the ARF3 protein repressed shoot regeneration by activating the SA signaling pathway. Furthermore, transcriptomic analysis combined with cis-regulatory motif screening identified candidate stress responses-related ARF3 target genes that function downstream of the CRY1 regulatory network. This supports a model in which CRY1 promotes shoot regeneration by restraining ARF3-activated stress responses. Collectively, these findings revealed a fundamental trade-off between defense responses and regenerative growth in plants, orchestrated by the CRY1-ARF3 module, highlighting a photoreceptor-driven mechanism that connects environmental light cues to developmental plasticity.

In our study, CRY1-mediated blue light signaling enhanced de novo shoot regeneration from callus in *Arabidopsis* and the spatiotemporal analysis of CRY1-GFP highlighted the key role of CRY1 throughout shoot regeneration (Fig 1). CRY1 suppressed ARF3 at two levels-by downregulating *ARF3* transcription and by a direct, light-independent interaction with ARF3. The spatiotemporal analysis of ARF3-GFP revealed dynamic expression patterns during shoot regeneration, initially showing weak signal in root tips and subsequently forming dot-like condensates in callus and shoot apical meristem regions. Proteasome inhibitor treatment significantly enhanced ARF3 accumulation, indicating that ARF3 protein underwent proteasomal degradation during the regeneration process. (Fig 2 and 4). Conversely, ARF3 repressed shoot regeneration by activating SA signaling that inhibit shoot regeneration. SA-related genes were abundant among the downregulated genes in *arf3*, and SA treatment suppressed shot regeneration specifically in *arf3* (Fig 3 and 5). Consistent with this mechanism, *cry1* mutant explants showed elevated ARF3 expression and induction of stress responses accompanied by reduced shoot regeneration, whereas *arf3* mutants exhibited enhanced shoot regeneration (Fig 7). Furthermore, transcriptomic analysis combined with cis-regulatory motif scanning identified candidate stress responses-related ARF3 target genes in the downstream of the CRY1 regulatory network (Fig 6), supporting the model that CRY1 promotes shoot regeneration by restraining ARF3-activated stress responses.

**Figure 7.**
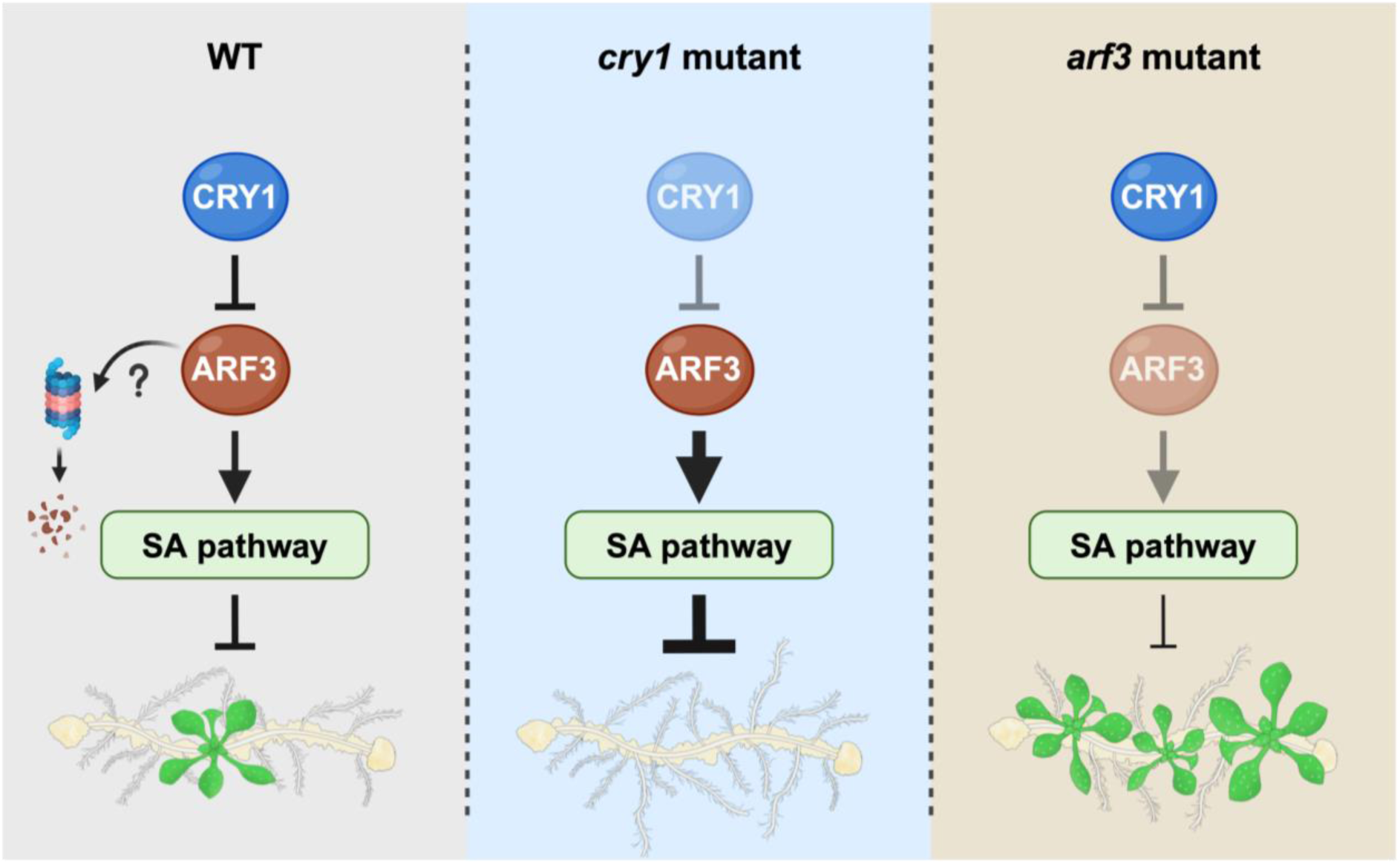
A model for depicting how CRY1 acts with ARF3 to regulate shoot regeneration from callus in Arabidopsis. CRY1-mediated blue light signaling promotes shoot regeneration by repressing the ARF3-activated SA pathway. CRY1 enhances shoot regeneration, primarily through repressing the transcription of *ARF3* and physical interaction with ARF3. Additionally, ARF3 may be degraded during shoot regeneration. Furthermore, ARF3 activates the SA pathway, which in turn represses shoot regeneration, highlighting the balance between the defense response and plant regeneration. In the absence of CRY1, ARF3 remains active, leading to enhanced SA pathway activation and reduced shoot regeneration. Conversely, in the *arf3* mutant, the suppression of shoot regeneration is alleviated due to reduced SA pathway activation. In the diagram, 50% transparent blue and brown ovals denote the absence of CRY1 and ARF3, respectively. Arrows and T-bars represent positive and negative regulation, respectively. 50% transparent arrows and T-bars denote the absence of regulation, while thicker and thinner arrows or T-bars represent stronger and weaker regulation, respectively.

Both BiFC and Co-IP protein interaction assays demonstrated that CRY1 interacts with ARF3 in a light-independent manner (Fig 4A and B). However, CRY1 has been previously shown to interact with ARF6/8 in a blue light-dependent manner (Mao et al., 2020). Recent studies have reported that more than half of the CRY-interacting proteins exhibit blue light-dependent change of interaction to CRYs (Qu et al., 2024). In contrast, several proteins interact with CRYs in a blue light-independent manner such as FKBP12-INTERACTING PROTEIN 37 (FIP37), TEOSINTE BRANCHED1-CYCLOIDEA-PCF 22 (TCP22) and MOS4-ASSOCIATED COMPLEX SUBUNITS 3A/3B (MAC3A/3B) (Wang et al., 2021; Mo et al., 2022; Jiang et al., 2023). These facts suggest that CRY complexes may retain biochemical and cellular activities even in darkness. Consistently, our shoot regeneration assays under red light showed that there was significant difference of shoot regeneration rate between WT and *cry1* under red light, which suggest that inactive CRY1 still has the activity to enhance shoot regeneration (Fig 1F and G). Collectively, these findings suggest that CRY1 could have suppressive effects on ARF3 under both light and dark conditions, implying that the regulatory mechanisms of CRY1 toward ARF3 differs from those previously described for ARF6 and ARF8.

Our spatiotemporal analysis of ARF3-GFP revealed that ARF3 formed condensates in callus tissue and SAM during shoot regeneration (Fig 4D), consistent with ARF3 patterns observed in Arabidopsis floral meristems and SAM (Zhang et al., 2022). Notably, OsARF3, the rice homolog of Arabidopsis ARF3 (Wang et al., 2007), similarly forms condensates (Lei et al., 2025). ARF3 has been characterized as a transcriptional repressor based on functional assays and its amino acid composition in the middle region (Tiwari et al., 2003; Guilfoyle and Hagen, 2007). However, our transcriptomic analysis revealed that during shoot regeneration, particularly at the C14 and C14S1 stages, the number of downregulated genes in *arf3* was distinctly higher than the number of upregulated genes, suggesting that ARF3 primarily functioned as a transcriptional activator at these stages. In contrast, at the C0 stage, the number of downregulated genes was lower than that of upregulated genes, indicating that ARF3 may also function as a transcriptional repressor at this initial stage (Fig 5A). This context-dependent dual function aligned with reports from flower development showing that ARF3 can function as both a transcriptional repressor and activator depending on developmental stage (Simonini et al., 2017; Zhang et al., 2018). Moreover, this stage-specific switch in ARF3 activity correlated with its condensate formation pattern: the proteasome inhibitor treatment revealed formation of ARF3-GFP condensates specifically after callus and shoot induction, but not in the root tip at C0 (Fig 4D), implying that condensate formation was an induction-dependent phenomenon associated with its functional activator state for shoot regeneration. We speculated that CRY1 might modulate ARF3’s phase behavior, thereby regulating ARF3 protein stability during shoot regeneration.

In our study, we investigated the molecular mechanism by which ARF3 was regulated by CRY1 at the post-translational level using protein interaction assays and proteasome inhibitor treatment (Fig 4). These experiments provided evidence that CRY1 directly interacted with ARF3 and may promote its degradation under both light and dark conditions, thereby limiting ARF3’s ability to activate stress-responsive pathways that suppress shoot regeneration. However, the mechanism by which CRY1 regulated ARF3 at the transcriptional level remained elusive (Fig 2F). Although CRY1 does not primarily function as a transcription factor but rather as a blue light receptor, it is possible that CRY1 may indirectly modulate *ARF3* expression. One possible scenario is that CRY1 represses transcriptional activity or gene expression of upstream regulators that activate *ARF3* expression. Collectively, these dual layers of regulation may cooperate to ensure that ARF3 levels are precisely controlled by CRY1 in response to blue light, allowing the plant to balance between shoot regeneration and stress responses.

It is widely known that plant stress responses including biotic responses by SA have trade-off relationships with plant growth and development (Florentin et al., 2013; Ikeuchi et al., 2019; Koo et al., 2024). Our study uncovered, for the first time, a pivotal role of stress response in determining plant regeneration in the context of light condition. Meanwhile, previous works also showed that stress conditions such as wounding and hypoxia treatment enhance callus formation (Koo et al., 2024). These findings suggest that plants possess sophisticated molecular mechanisms to determine whether to promote or suppress dedifferentiation or regeneration according to the fluctuating environmental conditions.

Understanding the CRY1-ARF3 module is crucial for plant adaptation and survival, as this interplay coordinates organ regeneration and stress responses with environmental conditions. Light and darkness distinctly induce shoot and root regeneration, respectively, ensuring that new organs develop appropriately for their context: shoots in aerial, light-rich environments and roots in dark, soil-rich environments. Even aerial organs exposed to shade may initiate root regeneration to secure adequate nutrient uptake. Moreover, since soil harbors a greater abundance of pathogens and viruses than air (Ruiz-Gil et al., 2020), plants employ robust root regeneration and stress response mechanisms in dark, soil-rich conditions. ARF3’s role in enhancing pathogen resistance in crops (Lei et al., 2025) further suggests that modulating the CRY1-ARF3 interaction can help balance growth and immunity in agricultural settings. Thus, the CRY1-ARF3 interplay and its downstream regulation of stress-responsive pathways are vital for optimizing plant growth and resilience, particularly in agriculture.

In conclusion, our work established a mechanistic link between blue light perception and regenerative development through the CRY1-ARF3 module. By directly repressing ARF3, CRY1 declined ARF3-driven activation of SA pathway, thereby shifting the balance in favor of shoot regeneration (Fig 7). This CRY1-ARF3 module illustrated how plants integrate environmental cues with hormonal signaling to orchestrate the balance between regenerative growth and defense responses.

## Materials and Methods

### Plant materials and growth conditions

All the *Arabidopsis thaliana* plants used in this study are in Col-0 ecotype background. The following mutants were used in the present study: *phyA-211* (Reed et al., 1994), *phyB-5* (Reed et al., 1993), *cry1-304* (Stoelzle et al., 2003), *cry2-1* (Guo et al., 1998), *phot1* (salk_146058), *phot2* (salk_142275), *ett-13/arf3* (salk_040513) (Pekker et al., 2005; Iwasaki et al., 2013), *arf4-1* (Pekker et al., 2005; Iwasaki et al., 2013). Mutants *phyA-211, phyB-5, cry1-304, and cry2-1* were kindly provided by Professor Tomonao Matsushita; *phot1* and *phot2* by Professor Shinichiro Inoue, and *ett-13/arf3* and *arf4-1* by Professor Chiyoko Machida.

The *cry1 arf3* mutant was prepared by genetic crossing, and its identity was verified by genotyping. Complementation lines pro*CRY1*:g*CRY1*-GFP/*cry1* and pro*ARF3*:g*ARF3*-GFP/*arf3* were generated by *Agrobacterium tumefaciens* (GV3101) Agrobacterium-mediated transformation (floral dip) and selected for Basta resistance. Plants were grown on soil or MGRL (Fujiwara et al., 1992) medium under a long-day (16 h light/8 h darkness) photoperiod at 22°C.

### Plasmid construction

To generate the constructs to express proteins in plants, the pB7WG2 vector (https://gatewayvectors.vib.be/) was digested with *Eco*32I and ligated to remove the cassette of the LR reaction. For complementation assays, the multi-cloning site (MCS)-GFP-pB7WG2 vector was generated by insertion of the GFP sequence into the *Spe*I and *Pst*I site using NEBuilder® HiFi (NEB). Next, the genomic sequences of *CRY1* and *ARF3* were amplified by PCR and inserted into the MCS-GFP-pB7WG2 vector through the *Kpn*I and *Eco*RV site using NEBuilder® HiFi. For BiFC assays, the 35S-MCS-nYFP-pB7WG2 vector was generated by insertion of the *35S* promoter sequence into the *Kpn*I and *Bam*HI site, and the n*YFP* sequence through the *Bam*HI and *Xba*I using NEBuilder® HiFi. The 35S-MCS-cYFP-pB7WG2 vector was generated by insertion of the c*YFP* sequence through the *Spe*I and *Pst*I in the place of n*YFP* using NEBuilder® HiFi. The coding sequences of *CRY1* and *ARF3* were amplified by PCR and inserted into the 35S-MCS-cYFP-pB7WG2 and 35S-nYFP-MCS-pB7WG2 vector, respectively, through the *Bam*HI-*Spe*I site and *Xba*I-*Spe*I site, respectively, using NEBuilder® HiFi. For co-immunoprecipitation assays, the *35S* promoter sequence was inserted into the MCS-GFP-pB7WG2 vector through the *Kpn*I and *Bam*HI site using NEBuilder® HiFi to generate the 35S-MCS-GFP-pB7WG2 vector. The 35S-Flag-MCS-pB7WG2 vector was generated by insertion of the 3x*Flag* sequence into the 35S-MCS-pB7WG2 vector through the *Bam*HI and *Xba*I using T4 DNA ligase (TaKaRa). The coding sequences of *CRY1* and *ARF3* were amplified by PCR and inserted into the 35S-MCS-GFP-pB7WG2 and 35S-Flag-MCS-pB7WG2 vector, respectively, through the *Bam*HI-*Spe*I site and *Xba*I-*Spe*I site, respectively, using NEBuilder® HiFi. The primers used in this study are listed in the Supplementary Table 1.

### Regeneration assays

Shoot regeneration assays followed a two-step culture as previously described (Li et al., 2024). After surface sterilization, the seeds were sown onto MGRL medium containing 1.75 mM sodium phosphate buffer (pH 5.8), 1.5 mM MgSO_4_, 2.0 mM Ca(NO_3_)_2_, 3.0 mM KNO_3_, 67 µM Na_2_EDTA, 8.6 µM FeSO_4_, 10.3 µM MnSO_4_, 30 µM H_3_BO_3_, 1.0 µM ZnSO_4_, 24 nM (NH_4_)_6_Mo_7_O_24_, 130 nM CoCl_12_, 1 µM CuSO_4_, 15 g L^−1^ sucrose (Sigma), and 15 g L^−1^ gellan gum (Wako). Following stratification for 2 days in darkness at 4°C, the MRGL plates with seeds were placed vertically in a growth chamber, and the seedlings were grown at 22°C for 6 days under a 16 h light/8 h darkness photoperiod with a light intensity of 15 µmol m^−2^ s^−1^. Next, root explants (1 cm from the root tip) were excised from 6-day-old seedlings and cultured on CIM containing Gamborg’s B5 salt mixture (Sigma) with 20 g L^−1^ glucose (Wako), 0.5 g L^−1^ MES (NACALAI TESQUE, INC.), 1 mL L^−1^ Gamborg’s vitamin solution (Sigma), 500 µg L^−1^ of 2,4-D (Sigma), 50 µg L^−1^ of kinetin (Sigma), and 8 g L^−1^ gellan gum (Wako). The pH was adjusted to 5.7 by adding 1.0 M KOH. Continuous light was used for callus induction with a light intensity of 15 µmol m^−2^ s^−1^ at 25°C. After culturing on CIM for 14 days, the explants were transferred onto SIM containing Gamborg’s B5 salt mixture (Sigma) with 10 g L^−1^ glucose (Wako), 0.5 g L^−1^ MES (NACALAI TESQUE, INC.), 1 mL L^−1^ Gamborg’s vitamin solution (Sigma), 2 µg mL^−1^ *trans*-zeatin (Tokyo Chemical Industry), 0.4 µg mL^−1^indole-3-butyric acid (Sigma), 1 µg mL^−1^ d-biotin (Sigma), and 8 g L^−1^ gellan gum (Wako). The pH was adjusted to 5.7 by adding 1.0 M KOH. Continuous light was used for shoot induction with a light intensity of 15 µmol m^−2^ s^−1^ at 25°C. Finally, after culturing on SIM for 14 days, shoot regeneration frequency per explant was statistically evaluated by microscopy observation (Nikon SMZ18). A structure exhibiting a visible apical meristem surrounded by three or more trichome-bearing leaves was counted as one shoot. Light-quality experiments used continuous white or red light from callus induction onward, light intensities were matched (15 µmol m⁻² s⁻¹). To examine SA effects during shoot induction, explants were incubated on SIM supplemented with 25 μM SA (Sigma).

### Confocal microscopy imaging

Confocal microscopy imaging was performed using an FV1200 confocal laser scanning microscope (Olympus) equipped with a UPLSAPO20X objective lens (N.A. = 0.75, W.D. = 0.6 mm). For the observation of CRY1-GFP and ARF3-GFP during the shoot regeneration process, root explants at stages C0, C14, and C14S1 were examined. GFP fluorescence was excited using a 473 nm laser line. Image acquisition and processing were conducted using FV10-ASW software ver. 4.2a (Olympus) and Fiji software (Schindelin et al., 2012).

### Transcriptome analysis

Root explants were collected at C0, C14, and C14S1 stages. Total RNA was isolated from the collected explants using the Maxwell RSC Plant RNA Kit (Promega). The integrity of purified RNA was assessed using a 2100 Bioanalyzer (Agilent). A total of 1000 ng RNA was used to construct a transcriptome library with NEBNext Ultra II RNA Library Prep Kit for Illumina (New England Biolabs) according to the supplier’s instruction. Libraries were qualified and quantified using 2100 Bioanalyzer (Agilent). Illumina NextSeq500 platform was used for paired-end sequencing. Three independent biological replicates were analyzed for each genotype.

Reads were quality-checked by FastQC v.0.11.5 (Andrews et al., 2010) and trimmed by Trimmomatic v.0.40 (Bolger et al., 2014). Clean reads were aligned to the Araport11 reference genome with STAR v.2.6.1d (Dobin et al., 2013), and gene-level counts were obtained using featureCounts v.2.0.6 (Liao et al., 2014). Differential expression was performed with EdgeR v.3.42.4 (Robinson et al., 2010), with significance thresholds |Log₂FC| ≥ 1 and FDR < 0.05. GO enrichment analysis was conducted using the ShinyGO v.0.80 (Ge et al., 2020).

### BiFC assays

All constructs for BiFC assays were transformed into *Agrobacterium tumefaciens* strain GV3101. Overnight cultures were resuspended in infiltration buffer containing 2.15 g L⁻¹ MS salt, 50 g L⁻¹ Sucrose. The pH of the buffer was adjusted to 5.7 by adding 1.0 M KOH. The resulting suspension was used for infiltration of *Nicotiana benthamiana* (tobacco) leaves (Sato et al., 2014). Plants were maintained for 4 days under long-day (16 h light/8 h dark) white light condition before imaging. The epidermal cells of the infected leaves were observed by an FV1200 confocal microscope (Olympus).

For light-independence tests, infiltrated leaves were kept in white light for 4 days and then moved to darkness for 1 day prior to imaging. YFP signals were recorded by confocal microscopy using matched settings across treatments. Negative controls included single-half constructs (CRY1-cYFP or nYFP-ARF3 alone) infiltrated with the complementary empty vector. This experiment was independently repeated twice with similar results.

### Co-immunoprecipitation (Co-IP) assays

Co-IP assays were performed as described previously (Ito et al., 2024). To perform Co-IP assays, *Agrobacterium tumefaciens* strain GV3101, carrying 35S:CRY1-GFP and 35S:Flag-ARF3 constructs was infiltrated into tobacco leaves, together with 35S:GFP as a negative control. After infiltration, plants were maintained for 5 days under white light or in darkness prior to harvest. Immunoprecipitation was performed using a μMACS GFP Isolation Kit (Miltenyi Biotec) according to the supplier’s instruction. Samples were grounded to fine powder in liquid nitrogen, and homogenized in μMACS lysis buffer (Miltenyi Biotec) with Protease Inhibitor Cocktail (Sigma-Aldrich). The lysate was filtered through Cell Strainer (VCS-40; AS ONE Corporation), mixed with μMACS Anti-GFP MicroBeads from the μMACS GFP Isolation Kit (Miltenyi Biotec), and incubated at 4°C for 3 h. The precipitated samples were washed with wash buffer and eluted with elution buffer. Anti-GFP antibody (ab290; Abcam) (1:2,500) and monoclonal Anti-FLAG® M2 antibody (F3165; Sigma-Aldrich) (1:1,000) were used as primary antibodies. Anti-IgG (H+L chain) (Rabbit) pAb-HRP (458, MBL) (1:5,000) and anti-mouse IgG (H+L) Antibody, HRP conjugate (W402B; Promega) (1:2,000), were used as the secondary antibodies.

### Bortezomib treatment

To examine ARF3 stability, root explants were transferred from solid MGRL, CIM or SIM media to liquid MGRL, CIM or SIM media containing mock control (0.1% DMSO) or bortezomib (40 μM) (FUJIFILM) dissolved in 0.1% DMSO for 8 h on a shaker (AS ONE) at 16 rpm, 27°C.

## Acknowledgements

We thank Tomonao Matsushita (Kyoto University, Japan) for *phyA-211, phyB-5, cry1-304, and cry2-1* seeds, Shinichiro Inoue (Saitama University, Japan) for *phot1* and *phot2* seeds, and Chiyoko Machida (Chubu University, Japan) for *ett-13/arf3* and *arf4-1* seeds. We also thank Sayoko Mibu at the University of Tokyo for technical assistance. This research was supported by grants from MEXT/JSPS KAKENHI (20H03297 and 22H00415 to S.M.), ASPIRE (JPMJAP2306 to S.M.), JST SPRING (JPMJSP2108 to M.L.), MEXT/JSPS KAKENHI (24K09498, 22K15137 and JP22H04925 (PAGS) to H.S.) and a fellowship grant from the Human Frontier Scientific Program (LT000162/2018-L to H.S.).

## Author contributions

**Min Li:** Conceptualization; Data curation; Resources; Formal analysis; Validation; Investigation; Visualization; Methodology; Funding acquisition; Writing – original draft; Writing – review and editing. **Hikaru Sato:** Conceptualization; Supervision; Data curation; Resources; Investigation; Methodology; Funding acquisition; Writing – review and editing. **Takuya Sakamoto:** Supervision. **Yayoi Inui:** Supervision. **Kazunari Yamamoto:** Data curation; Formal analysis; Investigation. **Ryosuke Makino:** Data curation; Formal analysis; Investigation. **Tomonao Matsushita:** Resources. **Sachihiro Matsunaga:** Conceptualization; Supervision; Project administration; Funding acquisition; Visualization; Writing – review and editing.

---

**Appendix Fig S1.**
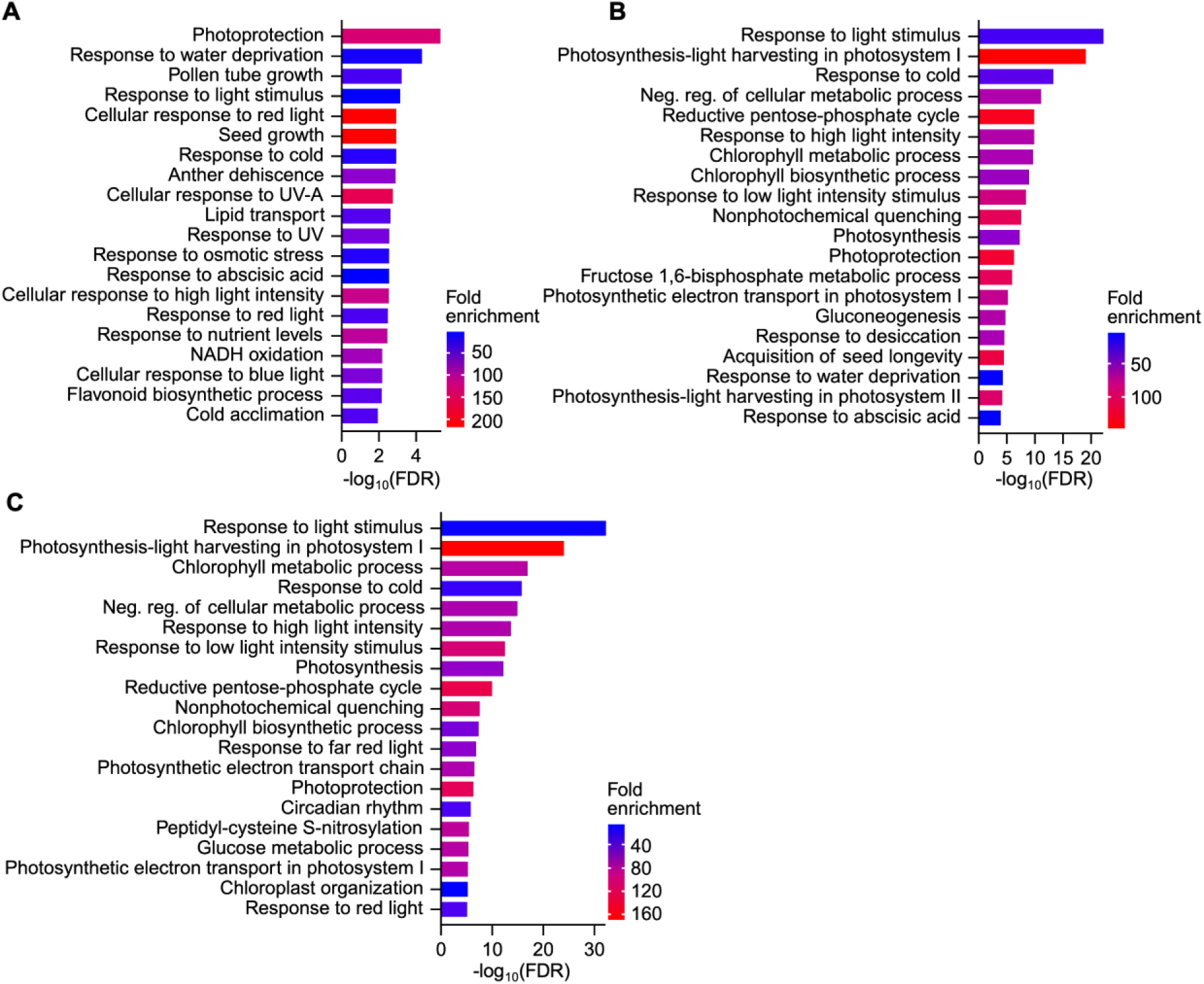
Transcriptomic analysis in *cry1-304* during de novo shoot regeneration. (A, B and C) Top 20 gene ontology (GO) terms enriched in genes downregulated in *cry1-304* compared with that in WT at C0 (A), C14 (B), and C14S1 (C) stages.

**Appendix Fig S2.**
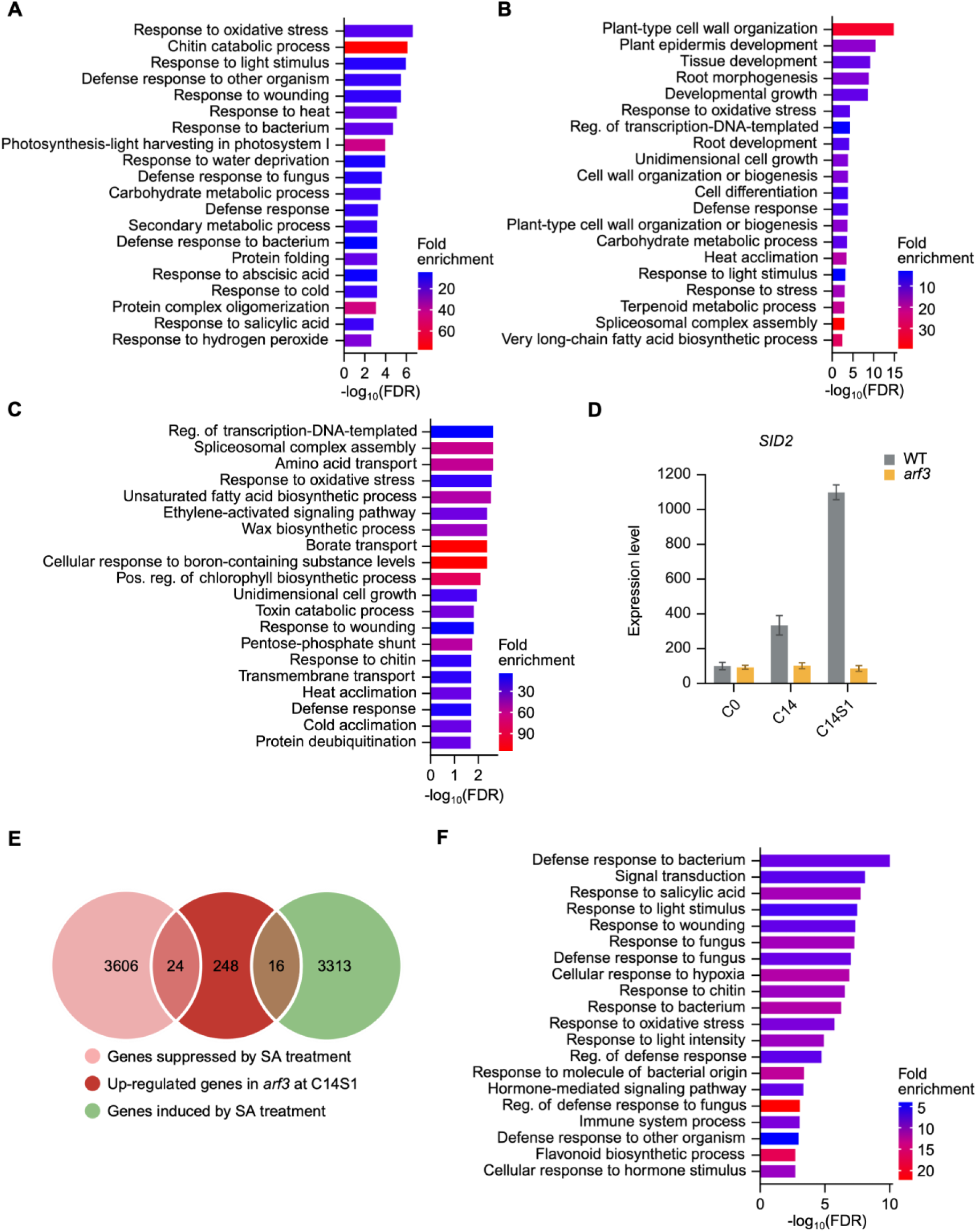
Transcriptomic analysis in *arf3* during de novo shoot regeneration. (A, B and C) Top 20 gene ontology (GO) terms enriched in genes upregulated in *arf3* compared with that in WT at C0 (A), C14 (B), and C14S1 (C) stages. (D) Expression pattern of *SID2* (Koo et al., 2024) in WT and *arf3* at C0, C14, and C14S1 stages. *SID2* was downregulated in *arf3* at the C14 and C14S1 stage. Relative expression was measured as normalized counts per million mapped reads, and the expression level of *SID2* in WT at C0 stage were set to 100. (E) Venn diagrams showing the overlap of the up-regulated genes in *arf3* at C14S1 stage (red) and the genes suppressed by salicylic acid (SA) treatment (pink) or the genes induced by SA treatment (green) (Zhang et al., 2020). (F) Top 20 gene ontology (GO) terms enriched in genes down-regulated in the *ett-3* loss-of-function mutant compared with that in WT during gynoecium development (Simonini et al, 2017).

## Tables and their legends

Supplementary Table 1. Primer list

Supplementary Table 2. Differential expression analysis between wild type and *cry1* mutant at C0

Supplementary Table 3. Differential expression analysis between wild type and *cry1* mutant at C14

Supplementary Table 4. Differential expression analysis between wild type and *cry1* mutant at C14S1

Supplementary Table 5. Differential expression analysis between wild type and *arf3* mutant at C0

Supplementary Table 6. Differential expression analysis between wild type and *arf3* mutant at C14

Supplementary Table 7. Differential expression analysis between wild type and *arf3* mutant at C14S1

Supplementary Table 8. 20 overlap genes between downregulated genes in *arf3* mutant and upregulated genes in *cry1* mutant at C14S1 stage

## Reference

Ahmad, M., and Cashmore, A.R. (1993). HY4 gene of A. thaliana encodes a protein with characteristics of a blue-light photoreceptor. Nature 366, 162–166.

Andrews, F.K., Segonds-Pichon, A., Biggins, L., Krueger, C., and Wingett, S. (2010). FastQC: a quality control tool for high throughput sequence data.

Atkinson, J.A., Rasmussen, A., Traini, R., Voß, U., Sturrock, C., Mooney, S.J., Wells, D.M., and Bennett, M.J. (2014). Branching out in roots: uncovering form, function, and regulation. Plant Physiol 166, 538–550.

Berrocal-Lobo, M., Molina, A., and Solano, R. (2002). Constitutive expression of ETHYLENE-RESPONSE-FACTOR1 in Arabidopsis confers resistance to several necrotrophic fungi. Plant J 29, 23–32.

Birnbaum, K.D., and Sánchez Alvarado, A. (2008). Slicing across kingdoms: regeneration in plants and animals. Cell 132, 697–710.

Boer, D.R., Freire-Rios, A., van den Berg, W.A., Saaki, T., Manfield, I.W., Kepinski, S., López-Vidrieo, I., Franco-Zorrilla, J.M., de Vries, S.C., Solano, R., Weijers, D., and Coll, M. (2014). Structural basis for DNA binding specificity by the auxin-dependent ARF transcription factors. Cell 156, 577–589.

Bolger, A.M., Lohse, M., and Usadel, B. (2014). Trimmomatic: a flexible trimmer for Illumina sequence data. Bioinformatics 30, 2114–2120.

Briggs, W.R., and Christie, J.M. (2002). Phototropins 1 and 2: versatile plant blue-light receptors. Trends Plant Sci 7, 204–210.

Bustamante, J.A., Miller, N.D., and Spalding, E.P. (2025). Separate sites of action for cry1 and phot1 blue-light receptors in the Arabidopsis hypocotyl. Curr Biol 35, 100–108.e104.

Cancé, C., Martin-Arevalillo, R., Boubekeur, K., and Dumas, R. (2022). Auxin response factors are keys to the many auxin doors. New Phytol 235, 402–419.

Chaves, I., Pokorny, R., Byrdin, M., Hoang, N., Ritz, T., Brettel, K., Essen, L.-O., Van Der Horst, G.T.J., Batschauer, A., and Ahmad, M. (2011). The cryptochromes: blue light photoreceptors in plants and animals. Annual review of plant biology 62, 335–364.

Chen, L., Cao, X., Li, Y., Liu, M., Liu, Y., Guan, Y., Ruan, J., Mao, Z., Wang, W., Yang, H.Q., and Guo, T. (2025). Photoexcited Cryptochrome 1 Interacts With SPCHLESS to Regulate Stomatal Development in Arabidopsis. Plant Cell Environ 48, 286–296.

Chen, Y., Ince, Y., Kawamura, A., Favero, D.S., Suzuki, T., and Sugimoto, K. (2024). ELONGATED HYPOCOTYL5-mediated light signaling promotes shoot regeneration in Arabidopsis thaliana. Plant Physiol 196, 2549–2564.

Cheng, Z.J., Wang, L., Sun, W., Zhang, Y., Zhou, C., Su, Y.H., Li, W., Sun, T.T., Zhao, X.Y., Li, X.G., Cheng, Y., Zhao, Y., Xie, Q., and Zhang, X.S. (2013). Pattern of auxin and cytokinin responses for shoot meristem induction results from the regulation of cytokinin biosynthesis by AUXIN RESPONSE FACTOR3. Plant Physiol 161, 240–251.

Cho, H., Ryu, H., Rho, S., Hill, K., Smith, S., Audenaert, D., Park, J., Han, S., Beeckman, T., Bennett, M.J., Hwang, D., De Smet, I., and Hwang, I. (2014). A secreted peptide acts on BIN2-mediated phosphorylation of ARFs to potentiate auxin response during lateral root development. Nat Cell Biol 16, 66–76.

Chory, J. (2010). Light signal transduction: an infinite spectrum of possibilities. Plant J 61, 982–991.

de Roij, M., Hernández García, J., Das, S., Borst, J.W., and Weijers, D. (2025). ARF degradation defines a deeply conserved step in auxin response. Nat Plants.

Dobin, A., Davis, C.A., Schlesinger, F., Drenkow, J., Zaleski, C., Jha, S., Batut, P., Chaisson, M., and Gingeras, T.R. (2013). STAR: ultrafast universal RNA-seq aligner. Bioinformatics 29, 15–21.

Florentin, A., Damri, M., and Grafi, G. (2013). Stress induces plant somatic cells to acquire some features of stem cells accompanied by selective chromatin reorganization. Dev Dyn 242, 1121–1133.

Franco-Zorrilla, J.M., López-Vidriero, I., Carrasco, J.L., Godoy, M., Vera, P., and Solano, R. (2014). DNA-binding specificities of plant transcription factors and their potential to define target genes. Proc Natl Acad Sci U S A 111, 2367–2372.

Fujiwara, T., Hirai, M.Y., Chino, M., Komeda, Y., and Naito, S. (1992). Effects of sulfur nutrition on expression of the soybean seed storage protein genes in transgenic petunia. Plant Physiol 99, 263–268.

Ge, S.X., Jung, D., and Yao, R. (2020). ShinyGO: a graphical gene-set enrichment tool for animals and plants. Bioinformatics 36, 2628–2629.

Guan, C., Wu, B., Yu, T., Wang, Q., Krogan, N.T., Liu, X., and Jiao, Y. (2017). Spatial auxin signaling controls leaf flattening in Arabidopsis. Current Biology 27, 2940–2950.

Guilfoyle, T.J., and Hagen, G. (2007). Auxin response factors. Curr Opin Plant Biol 10, 453–460.

Guo, H., Yang, H., Mockler, T.C., and Lin, C. (1998). Regulation of flowering time by Arabidopsis photoreceptors. Science 279, 1360–1363.

Hao, Y., Zeng, Z., Yuan, M., Li, H., Guo, S., Yang, Y., Jiang, S., Hawara, E., Li, J., Zhang, P., Wang, J., Xin, X., Ma, W., and Liu, H. (2025). The blue-light receptor CRY1 serves as a switch to balance photosynthesis and plant defense. Cell Host Microbe 33, 137–150.e136.

Hasnain, A., Naqvi, S.A.H., Ayesha, S.I., Khalid, F., Ellahi, M., Iqbal, S., Hassan, M.Z., Abbas, A., Adamski, R., Markowska, D., Baazeem, A., Mustafa, G., Moustafa, M., Hasan, M.E., and Abdelhamid, M.M.A. (2022). Plants. Front Plant Sci 13, 1009395.

Hernández-Coronado, M., Dias Araujo, P.C., Ip, P.L., Nunes, C.O., Rahni, R., Wudick, M.M., Lizzio, M.A., Feijó, J.A., and Birnbaum, K.D. (2022). Plant glutamate receptors mediate a bet-hedging strategy between regeneration and defense. Dev Cell 57, 451–465.e456.

Holtkotte, X., Ponnu, J., Ahmad, M., and Hoecker, U. (2017). The blue light-induced interaction of cryptochrome 1 with COP1 requires SPA proteins during Arabidopsis light signaling. PLoS Genet 13, e1007044.

Hu, Y., Chen, L., Wang, H., Zhang, L., Wang, F., and Yu, D. (2013). Arabidopsis transcription factor WRKY8 functions antagonistically with its interacting partner VQ9 to modulate salinity stress tolerance. Plant J 74, 730–745.

Ikeuchi, M., Ogawa, Y., Iwase, A., and Sugimoto, K. (2016). Plant regeneration: cellular origins and molecular mechanisms. Development 143, 1442–1451.

Ikeuchi, M., Favero, D.S., Sakamoto, Y., Iwase, A., Coleman, D., Rymen, B., and Sugimoto, K. (2019). Molecular Mechanisms of Plant Regeneration. Annu Rev Plant Biol 70, 377–406.

Ito, N., Sakamoto, T., Oko, Y., Sato, H., Hanamata, S., Sakamoto, Y., and Matsunaga, S. (2024). Nuclear pore complex proteins are involved in centromere distribution. iScience 27, 108855.

Iwasaki, M., Takahashi, H., Iwakawa, H., Nakagawa, A., Ishikawa, T., Tanaka, H., Matsumura, Y., Pekker, I., Eshed, Y., Vial-Pradel, S., Ito, T., Watanabe, Y., Ueno, Y., Fukazawa, H., Kojima, S., Machida, Y., and Machida, C. (2013). Dual regulation of ETTIN (ARF3) gene expression by AS1-AS2, which maintains the DNA methylation level, is involved in stabilization of leaf adaxial-abaxial partitioning in Arabidopsis. Development 140, 1958–1969.

Iwase, A., Harashima, H., Ikeuchi, M., Rymen, B., Ohnuma, M., Komaki, S., Morohashi, K., Kurata, T., Nakata, M., Ohme-Takagi, M., Grotewold, E., and Sugimoto, K. (2017). WIND1 Promotes Shoot Regeneration through Transcriptional Activation of. Plant Cell 29, 54–69.

Jiang, B., Zhong, Z., Su, J., Zhu, T., Yueh, T., Bragasin, J., Bu, V., Zhou, C., Lin, C., and Wang, X. (2023). Co-condensation with photoexcited cryptochromes facilitates MAC3A to positively control hypocotyl growth in. Sci Adv 9, eadh4048.

Jing, H., Korasick, D.A., Emenecker, R.J., Morffy, N., Wilkinson, E.G., Powers, S.K., and Strader, L.C. (2022). Regulation of AUXIN RESPONSE FACTOR condensation and nucleo-cytoplasmic partitioning. Nat Commun 13, 4015.

Koo, D., Lee, H.G., Bae, S.H., Lee, K., and Seo, P.J. (2024). Callus proliferation-induced hypoxic microenvironment decreases shoot regeneration competence in Arabidopsis. Mol Plant 17, 395–408.

Kuhn, A., Ramans Harborough, S., McLaughlin, H.M., Natarajan, B., Verstraeten, I., Friml, J., Kepinski, S., and Østergaard, L. (2020). Direct ETTIN-auxin interaction controls chromatin states in gynoecium development. Elife 9.

Lakehal, A., Chaabouni, S., Cavel, E., Le Hir, R., Ranjan, A., Raneshan, Z., Novák, O., Păcurar, D.I., Perrone, I., Jobert, F., Gutierrez, L., Bakò, L., and Bellini, C. (2019). A Molecular Framework for the Control of Adventitious Rooting by TIR1/AFB2-Aux/IAA-Dependent Auxin Signaling in Arabidopsis. Mol Plant 12, 1499–1514.

Lavy, M., Prigge, M.J., Tao, S., Shain, S., Kuo, A., Kirchsteiger, K., and Estelle, M. (2016). Constitutive auxin response in Physcomitrella reveals complex interactions between Aux/IAA and ARF proteins. Elife 5.

Lei, M.Q., He, R.R., Zhou, Y.F., Yang, L., Zhang, Z.F., Yuan, C., Zhao, W.L., Cheng, Y., Lian, J.P., Zhang, Y.C., Wang, W.T., Yu, Y., and Chen, Y.Q. (2025). The long noncoding RNA ALEX1 confers a functional phase state of ARF3 to enhance rice resistance to bacterial pathogens. Mol Plant 18, 114–129.

Li, M., Sato, H., and Matsunaga, S. (2024). Application of de novo shoot regeneration system from root explants of Arabidopsis thaliana. CYTOLOGIA 89, 83–84.

Liao, Y., Smyth, G.K., and Shi, W. (2014). featureCounts: an efficient general purpose program for assigning sequence reads to genomic features. Bioinformatics 30, 923–930.

Liu, H., Lin, R., and Deng, X.W. (2020). Photobiology: Light signal transduction and photomorphogenesis. J Integr Plant Biol 62, 1267–1269.

Liu, K., Li, Y., Chen, X., Li, L., Zhao, H., Wang, Y., and Han, S. (2018). ERF72 interacts with ARF6 and BZR1 to regulate hypocotyl elongation in Arabidopsis. J Exp Bot 69, 3933–3947.

Liu, S., Wang, Q., Zhong, M., Lin, G., Ye, M., Wang, Y., and Zhang, J. (2025a). The CRY1-COP1-HY5 axis mediates blue-light regulation of Arabidopsis thermotolerance. Plant Commun 6, 101264.

Liu, S., Wang, Q., Zhong, M., Lin, G., Ye, M., Wang, Y., and Zhang, J. (2025b). The CRY1-COP1-HY5 axis mediates blue-light regulation of Arabidopsis thermotolerance. Plant Commun, 101264.

Liu, S., Zhang, L., Gao, L., Chen, Z., Bie, Y., Zhao, Q., Zhang, S., Hu, X., Liu, Q., Wang, X., and Wang, Q. (2022). Differential photoregulation of the nuclear and cytoplasmic CRY1 in Arabidopsis. New Phytol 234, 1332–1346.

Luo, X., Xu, N., Huang, J., Gao, F., Zou, H., Boudsocq, M., Coaker, G., and Liu, J. (2017). A Lectin Receptor-Like Kinase Mediates Pattern-Triggered Salicylic Acid Signaling. Plant Physiol 174, 2501–2514.

Mao, Z., He, S., Xu, F., Wei, X., Jiang, L., Liu, Y., Wang, W., Li, T., Xu, P., Du, S., Li, L., Lian, H., Guo, T., and Yang, H.Q. (2020). Photoexcited CRY1 and phyB interact directly with ARF6 and ARF8 to regulate their DNA-binding activity and auxin-induced hypocotyl elongation in Arabidopsis. New Phytol 225, 848–865.

Martin-Arevalillo, R., Thévenon, E., Jégu, F., Vinos-Poyo, T., Vernoux, T., Parcy, F., and Dumas, R. (2019). Evolution of the Auxin Response Factors from charophyte ancestors. PLoS Genet 15, e1008400.

Maruyama, K., Todaka, D., Mizoi, J., Yoshida, T., Kidokoro, S., Matsukura, S., Takasaki, H., Sakurai, T., Yamamoto, Y.Y., Yoshiwara, K., Kojima, M., Sakakibara, H., Shinozaki, K., and Yamaguchi-Shinozaki, K. (2012). Identification of cis-acting promoter elements in cold- and dehydration-induced transcriptional pathways in Arabidopsis, rice, and soybean. DNA Res 19, 37–49.

Mo, W., Zhang, J., Zhang, L., Yang, Z., Yang, L., Yao, N., Xiao, Y., Li, T., Li, Y., Zhang, G., Bian, M., Du, X., and Zuo, Z. (2022). Arabidopsis cryptochrome 2 forms photobodies with TCP22 under blue light and regulates the circadian clock. Nat Commun 13, 2631.

Motte, H., Vereecke, D., Geelen, D., and Werbrouck, S. (2014). The molecular path to in vitro shoot regeneration. Biotechnol Adv 32, 107–121.

Mutte, S.K., Kato, H., Rothfels, C., Melkonian, M., Wong, G.K., and Weijers, D. (2018). Origin and evolution of the nuclear auxin response system. Elife 7.

Nameth, B., Dinka, S.J., Chatfield, S.P., Morris, A., English, J., Lewis, D., Oro, R., and Raizada, M.N. (2013). The shoot regeneration capacity of excised Arabidopsis cotyledons is established during the initial hours after injury and is modulated by a complex genetic network of light signalling. Plant Cell Environ 36, 68–86.

Ohgishi, M., Saji, K., Okada, K., and Sakai, T. (2004). Functional analysis of each blue light receptor, cry1, cry2, phot1, and phot2, by using combinatorial multiple mutants in Arabidopsis. Proc Natl Acad Sci U S A 101, 2223–2228.

Park, O.S., Bae, S.H., Kim, S.G., and Seo, P.J. (2019). JA-pretreated hypocotyl explants potentiate. Plant Signal Behav 14, 1618180.

Pasternak, T., Groot, E.P., Kazantsev, F.V., Teale, W., Omelyanchuk, N., Kovrizhnykh, V., Palme, K., and Mironova, V.V. (2019). Salicylic Acid Affects Root Meristem Patterning via Auxin Distribution in a Concentration-Dependent Manner. Plant Physiol 180, 1725–1739.

Pedmale, U.V., Huang, S.C., Zander, M., Cole, B.J., Hetzel, J., Ljung, K., Reis, P.A.B., Sridevi, P., Nito, K., Nery, J.R., Ecker, J.R., and Chory, J. (2016). Cryptochromes Interact Directly with PIFs to Control Plant Growth in Limiting Blue Light. Cell 164, 233–245.

Pekker, I., Alvarez, J.P., and Eshed, Y. (2005). Auxin response factors mediate Arabidopsis organ asymmetry via modulation of KANADI activity. The Plant Cell 17, 2899–2910.

Ponnu, J., and Hoecker, U. (2022). Signaling Mechanisms by Arabidopsis Cryptochromes. Front Plant Sci 13, 844714.

Qu, G.P., Jiang, B., and Lin, C. (2024). The dual-action mechanism of Arabidopsis cryptochromes. J Integr Plant Biol 66, 883–896.

Quail, P.H. (2002). Phytochrome photosensory signalling networks. Nat Rev Mol Cell Biol 3, 85–93.

Reed, J.W., Nagpal, P., Poole, D.S., Furuya, M., and Chory, J. (1993). Mutations in the gene for the red/far-red light receptor phytochrome B alter cell elongation and physiological responses throughout Arabidopsis development. Plant Cell 5, 147–157.

Reed, J.W., Nagatani, A., Elich, T.D., Fagan, M., and Chory, J. (1994). Phytochrome A and Phytochrome B Have Overlapping but Distinct Functions in Arabidopsis Development. Plant Physiol 104, 1139–1149.

Robinson, M.D., McCarthy, D.J., and Smyth, G.K. (2010). edgeR: a Bioconductor package for differential expression analysis of digital gene expression data. Bioinformatics 26, 139–140.

Ruiz-Gil, T., Acuña, J.J., Fujiyoshi, S., Tanaka, D., Noda, J., Maruyama, F., and Jorquera, M.A. (2020). Airborne bacterial communities of outdoor environments and their associated influencing factors. Environ Int 145, 106156.

Salmon, J., Ramos, J., and Callis, J. (2008). Degradation of the auxin response factor ARF1. Plant J 54, 118–128.

Sato, H., Mizoi, J., Tanaka, H., Maruyama, K., Qin, F., Osakabe, Y., Morimoto, K., Ohori, T., Kusakabe, K., and Nagata, M. (2014). Arabidopsis DPB3-1, a DREB2A interactor, specifically enhances heat stress-induced gene expression by forming a heat stress-specific transcriptional complex with NF-Y subunits. The Plant Cell 26, 4954–4973.

Schindelin, J., Arganda-Carreras, I., Frise, E., Kaynig, V., Longair, M., Pietzsch, T., Preibisch, S., Rueden, C., Saalfeld, S., Schmid, B., Tinevez, J.Y., White, D.J., Hartenstein, V., Eliceiri, K., Tomancak, P., and Cardona, A. (2012). Fiji: an open-source platform for biological-image analysis. Nat Methods 9, 676–682.

Schoenaers, S., Balcerowicz, D., Breen, G., Hill, K., Zdanio, M., Mouille, G., Holman, T.J., Oh, J., Wilson, M.H., Nikonorova, N., Vu, L.D., De Smet, I., Swarup, R., De Vos, W.H., Pintelon, I., Adriaensen, D., Grierson, C., Bennett, M.J., and Vissenberg, K. (2018). The Auxin-Regulated CrRLK1L Kinase ERULUS Controls Cell Wall Composition during Root Hair Tip Growth. Curr Biol 28, 722–732.e726.

Shukla, V., Lombardi, L., Iacopino, S., Pencik, A., Novak, O., Perata, P., Giuntoli, B., and Licausi, F. (2019). Endogenous Hypoxia in Lateral Root Primordia Controls Root Architecture by Antagonizing Auxin Signaling in Arabidopsis. Mol Plant 12, 538–551.

Simonini, S., Bencivenga, S., Trick, M., and Østergaard, L. (2017). Auxin-Induced Modulation of ETTIN Activity Orchestrates Gene Expression in Arabidopsis. Plant Cell 29, 1864–1882.

Simonini, S., Deb, J., Moubayidin, L., Stephenson, P., Valluru, M., Freire-Rios, A., Sorefan, K., Weijers, D., Friml, J., and Østergaard, L. (2016). A noncanonical auxin-sensing mechanism is required for organ morphogenesis in Arabidopsis. Genes Dev 30, 2286–2296.

Stoelzle, S., Kagawa, T., Wada, M., Hedrich, R., and Dietrich, P. (2003). Blue light activates calcium-permeable channels in Arabidopsis mesophyll cells via the phototropin signaling pathway. Proceedings of the National Academy of Sciences 100, 1456–1461.

Sugimoto, K., Temman, H., Kadokura, S., and Matsunaga, S. (2019). To regenerate or not to regenerate: factors that drive plant regeneration. Curr Opin Plant Biol 47, 138–150.

Sukiran, N.L., Ma, J.C., Ma, H., and Su, Z. (2019). ANAC019 is required for recovery of reproductive development under drought stress in Arabidopsis. Plant Mol Biol 99, 161–174.

Tiwari, S.B., Hagen, G., and Guilfoyle, T. (2003). The roles of auxin response factor domains in auxin-responsive transcription. Plant Cell 15, 533–543.

Ulmasov, T., Hagen, G., and Guilfoyle, T.J. (1997). ARF1, a transcription factor that binds to auxin response elements. Science 276, 1865–1868.

Valvekens, D., Van Montagu, M., and Van Lijsebettens, M. (1988). Agrobacterium tumefaciens-mediated transformation of Arabidopsis thaliana root explants by using kanamycin selection. Proc Natl Acad Sci U S A 85, 5536–5540.

Wang, D., Pei, K., Fu, Y., Sun, Z., Li, S., Liu, H., Tang, K., Han, B., and Tao, Y. (2007). Genome-wide analysis of the auxin response factors (ARF) gene family in rice (Oryza sativa). Gene 394, 13–24.

Wang, F., Han, T., and Jeffrey Chen, Z. (2024). Circadian and photoperiodic regulation of the vegetative to reproductive transition in plants. Commun Biol 7, 579.

Wang, Q., and Lin, C. (2020). Mechanisms of Cryptochrome-Mediated Photoresponses in Plants. Annu Rev Plant Biol 71, 103–129.

Wang, X., Jiang, B., Gu, L., Chen, Y., Mora, M., Zhu, M., Noory, E., Wang, Q., and Lin, C. (2021). A photoregulatory mechanism of the circadian clock in Arabidopsis. Nat Plants 7, 1397–1408.

Wei, X., Ding, Y., Wang, Y., Li, F., and Ge, X. (2020). Early Low-Fluence Red Light or Darkness Modulates the Shoot Regeneration Capacity of Excised. Plants (Basel) 9.

Wei, Y., Wang, S., and Yu, D. (2023). The Role of Light Quality in Regulating Early Seedling Development. Plants (Basel) 12.

Xu, J., Hofhuis, H., Heidstra, R., Sauer, M., Friml, J., and Scheres, B. (2006). A molecular framework for plant regeneration. Science 311, 385–388.

Xu, X., Kathare, P.K., Pham, V.N., Bu, Q., Nguyen, A., and Huq, E. (2017). Reciprocal proteasome-mediated degradation of PIFs and HFR1 underlies photomorphogenic development in. Development 144, 1831–1840.

Yang, Z., Liu, B., Su, J., Liao, J., Lin, C., and Oka, Y. (2017). Cryptochromes Orchestrate Transcription Regulation of Diverse Blue Light Responses in Plants. Photochem Photobiol 93, 112–127.

Yu, X., Liu, H., Klejnot, J., and Lin, C. (2010). The Cryptochrome Blue Light Receptors. Arabidopsis Book 8, e0135.

Zeng, J., Wang, Q., Lin, J., Deng, K., Zhao, X., Tang, D., and Liu, X. (2010). Arabidopsis cryptochrome-1 restrains lateral roots growth by inhibiting auxin transport. J Plant Physiol 167, 670–673.

Zhai, N., and Xu, L. (2021). Pluripotency acquisition in the middle cell layer of callus is required for organ regeneration. Nat Plants 7, 1453–1460.

Zhang, K., Zhang, H., Pan, Y., Guo, L., Tian, S., Wei, J., Fu, Y., Wang, C., Qu, P., and Liu, L. (2022). Cell-and non-cell-autonomous ARF3 coordinates meristem proliferation and organ patterning in Arabidopsis. bioRxiv, 2022-2001.

Zhang, K., Wang, R., Zi, H., Li, Y., Cao, X., Li, D., Guo, L., Tong, J., Pan, Y., Jiao, Y., Liu, R., Xiao, L., and Liu, X. (2018). AUXIN RESPONSE FACTOR3 Regulates Floral Meristem Determinacy by Repressing Cytokinin Biosynthesis and Signaling. Plant Cell 30, 324–346.

Zhang, M.M., Zhang, H.K., Zhai, J.F., Zhang, X.S., Sang, Y.L., and Cheng, Z.J. (2021). ARF4 regulates shoot regeneration through coordination with ARF5 and IAA12. Plant Cell Rep 40, 315–325.

Zhang, N., Zhou, S., Yang, D., and Fan, Z. (2020). Revealing Shared and Distinct Genes Responding to JA and SA Signaling in Arabidopsis by Meta-Analysis. Front Plant Sci 11, 908.

Zhang, T.Q., Lian, H., Zhou, C.M., Xu, L., Jiao, Y., and Wang, J.W. (2017). A Two-Step Model for de Novo Activation of. Plant Cell 29, 1073–1087.

Zhou, W., Lozano-Torres, J.L., Blilou, I., Zhang, X., Zhai, Q., Smant, G., Li, C., and Scheres, B. (2019). A Jasmonate Signaling Network Activates Root Stem Cells and Promotes Regeneration. Cell 177, 942–956.e914.

Zhou, Y., Liu, P., Tang, Y., Liu, J., Zhuang, Y., Li, X., Xu, K., Zhou, Z., Li, J., He, G., Deng, X.W., and Yang, L. (2024). NPR1 promotes blue light-induced plant photomorphogenesis by ubiquitinating and degrading PIF4. Proc Natl Acad Sci U S A 121, e2412755121.

